# Modulation of somatosensory signal transmission in the primate cuneate nucleus during voluntary hand movement

**DOI:** 10.1101/2023.05.09.540100

**Authors:** Shinji Kubota, Chika Sasaki, Satomi Kikuta, Sho Ito, Hiroaki Gomi, Tomomichi Oya, Kazuhiko Seki

**Affiliations:** Department of Neurophysiology, National Center of Neurology and Psychiatry, Tokyo, Japan; NTT Communication Science Laboratories, Nippon Telegraph and Telephone Co., Kanagawa, Japan

**Keywords:** Cuneate nucleus, Somatosensory gating, Voluntary movement, Primate, Proprioceptive afferent, Cutaneous afferent

## Abstract

Successful extraction of tactile and kinematic information is crucial for appropriate motor actions in daily life. To achieve this objective, sensory signals should be effectively regulated during motor actions. This regulation is understood both empirically and conceptually, but it is not well known where and how it is implemented in the central nervous system. Here, we show that, during voluntary movement, sensory signals are already attenuated in the primate cuneate nucleus, an early processing site in the ascending lemniscus pathway. The degree of suppression was comparable with the one reported in the cortex, suggesting that psychological attenuation of somatosensation could be ascribed to the cuneate. The results also revealed that this sensory attenuation was of descending origin, suggesting that cortical sensory prediction signals could regulate cuneate sensory transmission for extracting meaningful, and attenuate unnecessary, signals for movement regulation. This recurrent sensory modulation mechanism between cortical and subcortical areas may generalize to other sensory modalities and cognitive processes.

## Introduction

Manual dexterity, the ability to use your hand in a skillful way, is achieved by using somatosensory feedback signals.^1^ These signals, however, can also hinder dexterous movement when they are noisy or irrelevant to the ongoing action.^2, 3^ Thus, somatosensory signals from the hand must be regulated appropriately when they are integrated into the motor command.

During voluntary actions, somatosensory signals in the central nervous system (CNS) are selectively attenuated.^4–7^ This modulation, known as sensory attenuation or sensory gating,^8^ may reflect neuronal processes that attenuate the predicted, self-generated sensory feedback (reafference) in the somatosensory signal, thus isolating the unpredicted component of this signal.^7^ The latter component is important for error-based motor learning,^9^ and for extracting information about the external world via active exploration.^10^

While the theoretical significance of sensory gain modulation is established,^11^ its neural implementation remains unclear. Descending signals from cortical motor centers may be responsible for this phenomenon, because they support sensory prediction in the forward model (in the form of efference copy signals, ^12–14^ for example). The CNS contains several structures where somatosensory signals and descending motor commands converge,^15–18^ such as the cervical spinal cord. We previously characterized the neural mechanisms of sensory gain modulation in this structure,^19^ which receives somatosensory signals from the hand and the arm. We found that tactile and proprioceptive afferent signals were modulated during hand movement in behaving monkeys,^20^ and that the modulation could be generated via presynaptic inhibition.^19^ While the recorded neuron may be responsible for the central representation of somatosensory information along the postsynaptic dorsal column pathway,^21^ it may also regulate motor output along the spinal reflex pathway. It is currently unknown to what extent activity in the spinal cord affects the perception of somatosensory signals in the somatosensory ascending pathway and at the level of potential sensorimotor targets (e.g., sensorimotor cortex).

In parallel with the spinal cord, the cuneate nucleus (CN) in the medulla oblongata represents one of the first processing sites in the ascending lemniscus pathway. Neurons in the CN receive ascending somatosensory information from the upper body, and are exclusively involved in the ascending sensory pathway to the sensorimotor cortex or cerebrum. Furthermore, they receive descending signals from cortical and/or subcortical motor areas.^22–26^ This convergent projection from both afferent and efferent pathways^27–28^ suggests that sensory attenuation may occur at this site, thereby shaping cortical somatosensory perception in the lemniscus pathway and motor learning process in the cerebrum. This ascending pathway is crucial for voluntary hand movement, because perturbing the associated circuits results in abnormal manual behavior in transgenic mice.^29^ However, few previous studies have investigated the neural mechanism of sensory attenuation in the CN during voluntary movements,^30^ and no study existed in the primate CN, due to the technical challenge in recording neural signals from the deep location of this structure in the CNS using conventional experimental methods.^31–32^

To address this issue, we first established a method for precise electrode insertion into the subarea of macaque CN where neurons receive somatosensory afferent input from the hand. We then recorded the activity of CN neurons and characterized their response probability to electrical stimuli applied to cutaneous and proprioceptive afferents in monkeys performing wrist movements. We found an attenuation of cutaneous and proprioceptive ascending signal transmission during and even before wrist movement. The specific characteristics of this attenuation suggest that it is driven by both central motor commands and peripheral afferent feedback. These results indicate that the gain of somatosensory afferent signals is modulated in the CN for the purpose of providing higher sensory centers with low-noise signals from redundant sensory inputs.^33–35^

## Results

### Recording neural activity from primate cuneate nucleus

The CN is composed of two discrete substructures: the main cuneate and the external cuneate. The main cuneate receives cutaneous and proprioceptive signals from the upper limb and sends to them the thalamus; a pathway called the medial-lemniscus pathway, while the external cuneate receives proprioceptive signals and send them mainly to the cerebellum.^36–37^ In this study, we focused on the main cuneate. In primates, the main cuneate consists of five sub-areas distributed along the rostro-caudal direction (“rostral,” “cluster,” “shell,” “triangularis,” and “caudal” areas), each having unique input-output characteristics.^37^ For example, the “cluster” area receives cutaneous afferent inputs from the forelimb and hand, and projects to the ventral posterior lateral nucleus (VPL) of the thalamus. In addition to cutaneous afferent inputs, triangularis and rostral areas receive afferent inputs from muscles and joints, and project to the border of the VPL or to the ventral lateral nucleus of the thalamus.^36^ These region-specific input-output profiles prompted us to develop a method for probing each sub-area selectively via targeted insertion of the recording electrode (see below).

Before inserting the recording electrode into the brain, we identified the 3D coordinates of CN with an MRI scan. We then calculated optimal position, angle, and depth of insertion for each electrode penetration to target the CN by relying on aligned images from both CT and MRI scans^38^ (Figures1A and S1A) carried out in all monkeys. During penetrations, electrode travelled from the cortical surface to the CN over a distance of 4.35±0.14 cm in monkey KI, and 4.41±0.81 cm in monkey KE. We confirmed successful placement of the electrode inside the CN by examining neuronal responses to natural stimulation of the ipsilateral hindlimb, face, and forelimb (Figure 1B). We repeated this procedure for several penetrations at the beginning of each recording session and for each monkey, and we created a cortical surface map representing each penetration point and the targeted sub-area within the CN. We expanded this map by marking locations responsive to peripheral nerve stimulation during subsequent penetrations (Figure S1B). We applied electrical stimuli to the nerve cuff electrodes implanted around the superficial^19, 39^ (SR) and deep branch ^20^ (DR) of radial nerves (Figure 1C), and measured the size of evoked responses in the ipsilateral CN (see Figures S2 A and B for a detailed example).

**Figure 1:**
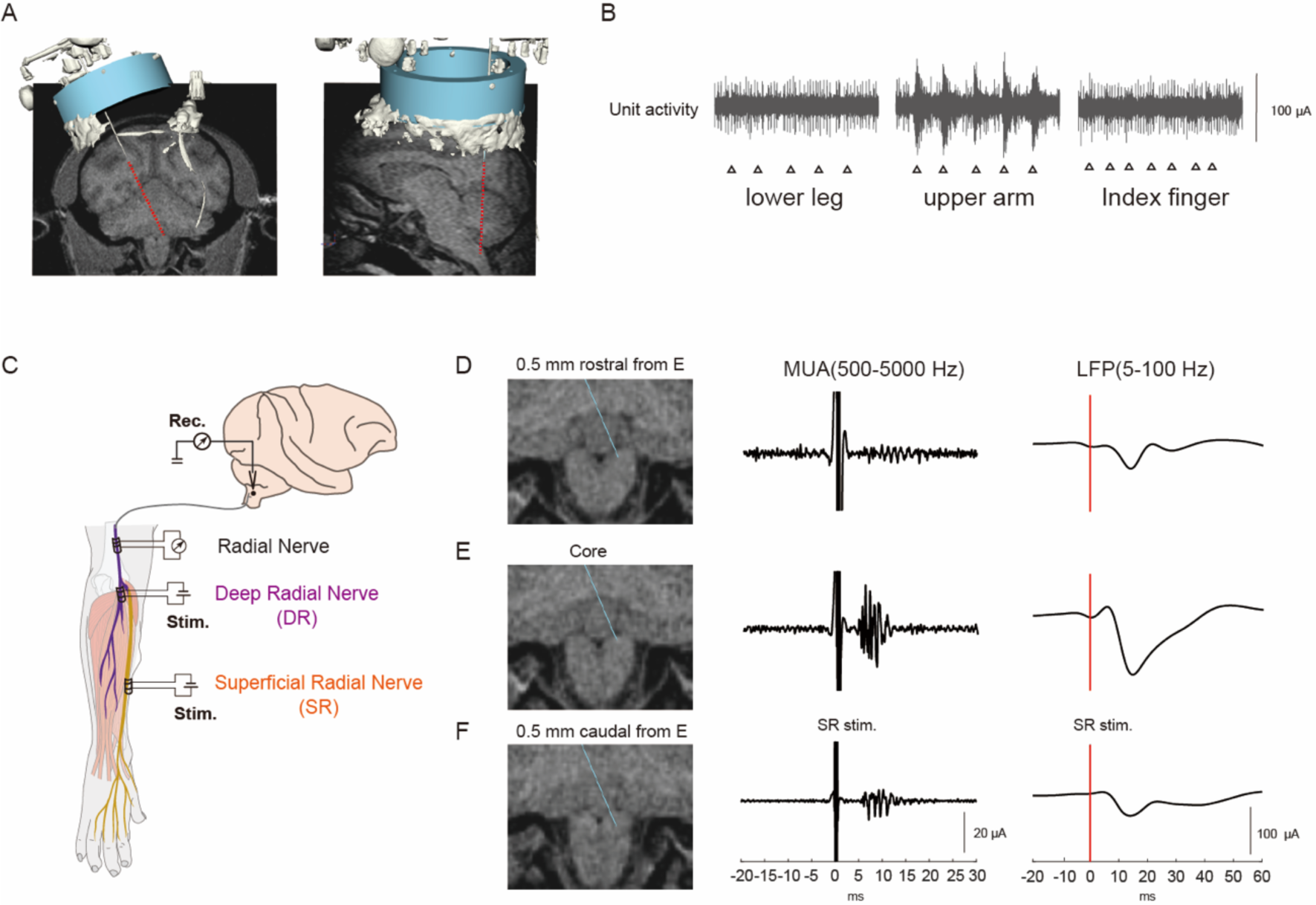
Recording Setup. **A,** Electrode trajectory to the cuneate nucleus (CN). MRI image with recording chamber and electrode reconstructed from aligned CT data showing coronal (**Left**) and sagittal (**Right**) views. The red dotted line indicates the estimated electrode trajectory. These composite images of CT and MRI were provided by Cicerone software.^38^ **B,** Responses of a CN neuron to light touches recorded at the target location. The lower leg (**Left**), upper arm (**Middle**), and index finger (**Right**) were stimulated by the experimenter’s hand (triangles). This neuron has a tactile receptive field on the upper arm. **C,** Peripheral nerve stimulation. Three nerve cuff electrodes were chronically implanted in the area of the superficial radial branch (SR), deep radial branch (DR), and radial stem (R) nerve of the right arm. The SR branch exclusively innervates a patch of skin on the dorsal, radial aspect of the hand. The DR branch mostly consists of muscle afferents from extensor muscles in the forearm. The SR and DR cuffs were used for stimulation (stim) and a proximal R cuff was used for recording elicited volleys. **D-F**, **Left column**; Coronal views of the recording sites in the SR response area of the CN, which showed clear SR-evoked responses. **Middle and Right columns**; Typical examples of SR-evoked multi-unit activities (MUA, filtered 500–5000 Hz, middle) and local field potentials (LFP, filtered 5– 100 Hz, right) recorded in each location. SR-evoked MUA and LFP of the CN were observed in each location, but the most prominent of which was recorded in the core point (E). The 0 ms in the middle and right panels indicates the timing of nerve stimulation (in this example, SR stimulation). The neural responses of CN were recorded extracellularly using single or multi-channel electrodes (8- or 16-channel S probe, Plexon Inc.).

Figure 1D–F shows a set of recordings from the SR-response area of the CN. The presence of dominant SR-evoked local field potentials (LFPs) and multi-unit activities (D–F) is consistent with the earlier characterization of this cluster area as exclusively receiving dense inputs from tactile afferents of the forearm.^27, 40^ The size of the evoked LFPs is larger at the core sites (E; peak amplitude of 125.10±50.45 µV) compared with rostral (D; 48.44±9.64 µV) and caudal sites (F; 48.70±39.38 µV). Moreover, these areas were clearly segregated within the region that showed tactile responses to a light touch of the leg (Figure S1B). These results confirm that our method can selectively record neuronal responses from targeted sub-areas within the CN.

### Attenuation of somatosensory input to the cuneate nucleus, and its modal specificity

Having identified the location of the CN and associated sub-areas using the focal targeting method detailed above, we next aimed to discover whether the transmission of afferent input to the CN is modulated during the execution of voluntary movement. For this aim, we trained two macaque monkeys to perform a wrist flexion-extension movement using a spring-loaded manipulandum (Figure 2A), and concomitantly recorded neural activity from the CN. Representative examples of single flexion/extension trials show that the amplitude of SR-evoked LFPs (SR-LFPs) started decreasing at around the onset time of dynamic wrist movement, for both flexion and extension (Figure 2B, top and bottom). For these examples, we found that the area of the SR-LFPs was significantly reduced during active movement [by 56.16% for flexion and 78.16% for extension, both significant at p<0.001 (flexion) and p=0.002 (extension) on a one-sample *t*-test], and during the hold period in flexion trials (80.02%; p=0.002) (Figure 2C). Figure 2D shows an example of DR-evoked LFPs around the time of movement onset. Again, we found that the DR-LFPs was greatly suppressed during movement and during the hold epoch, for both flexion and extension (Figure 2D, top and bottom). The area of the DR-LFPs in this example was significantly reduced during active movement [73.43 % for flexion (p=0.004) and 65.62% for extension (p<0.001)], and during the hold period in flexion trials (67.43%; p<0.001) (Figure 2E). Importantly, these reductions in the size of evoked LFPs cannot be attributed to the instability of the recording or stimulating electrodes implanted in the radial nerve, because electrical stimulation applied through both SR and DR cuff electrodes activated radial nerve afferents consistently, even during dynamic hand movement (Figure S2C–F, p=0.71 on a one-way ANOVA with post hoc Sidak’s test).

**Figure 2:**
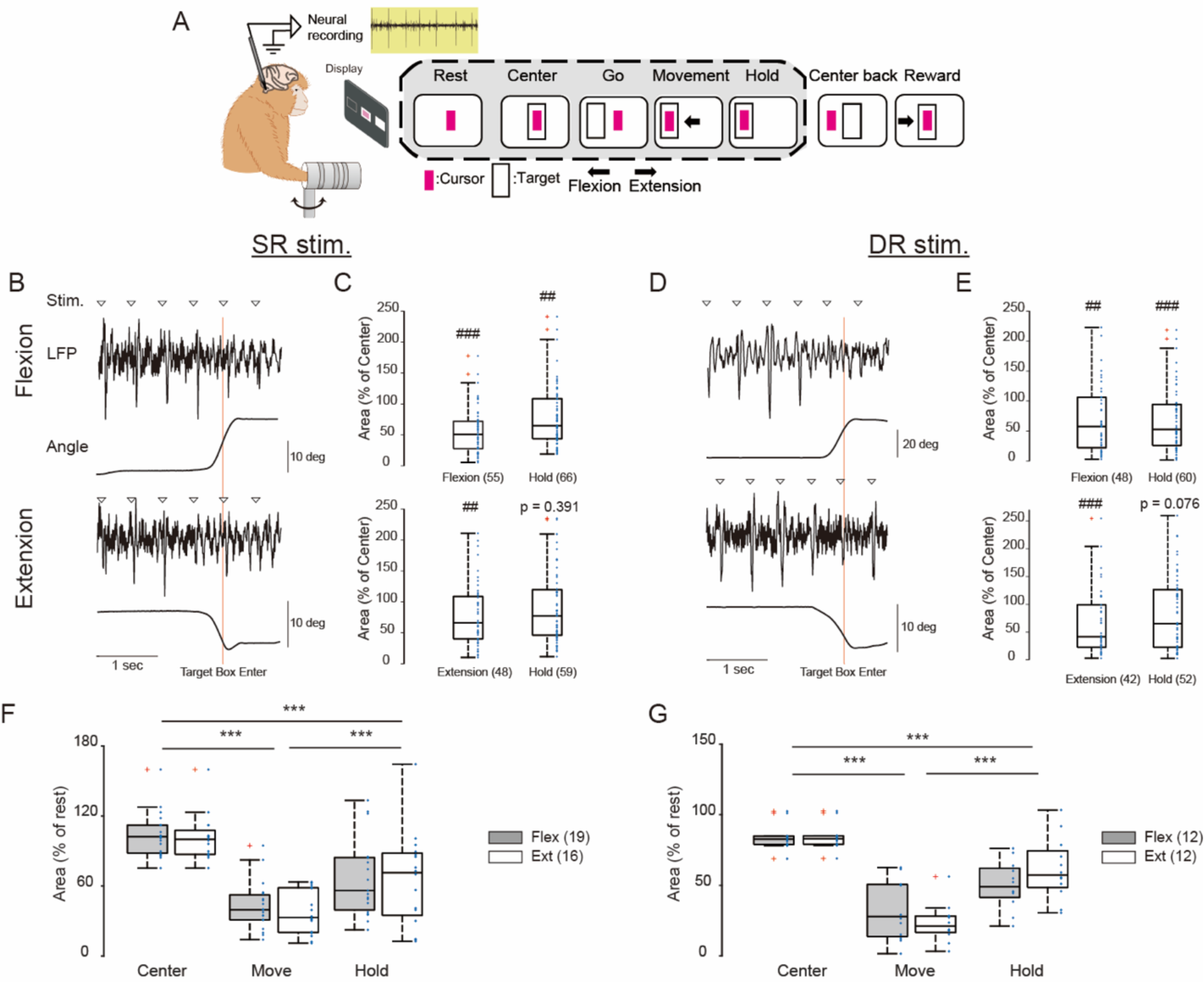
Modulation of evoked field potentials in the cuneate nucleus during a motor task. **A,** Task sequences. Monkeys performed a wrist flexion-extension movement with the aid of visual feedback of wrist position using a spring-loaded manipulandum. A red-filled square represents the moving cursor, the solid empty square represents the central and peripheral target. Black arrows indicate movement direction. Epochs marked in gray represent those analyzed in the present study. An illustration of single flexion trials. **B**, Typical examples of SR-evoked local field potential (LFP) in the cuneate nucleus (CN) during a single flexion (**Top**) and extension trial (**Bottom**). Triangles indicate the timing of the nerve stimuli (stim.). The red line illustrates the timing of the flexion or extension target box entry. **C,** Epoch-dependent modulation of the area of SR-evoked potentials shown in B. Data in both flexion and extension trials were averaged separately. Blue symbols denote data for the individual responses. Red crosses represent outlier data. Note that the reduction of SR-evoked potentials occurs during active movement regardless of movement direction. ## p < 0.01, ### p < 0.001 represents a significant difference compared with the center epoch (one sample t-test). **D** and **E,** Same as B and C, except that the DR was stimulated during a task. Note that DR-evoked LFP was also suppressed during active movement regardless of movement direction. ## p < 0.01, ### p < 0.001 represents a significant difference compared with the center epoch (one sample t-test). **F and G,** Boxplots showing the areas of SR-(**F**) and DR- **(G)** evoked potentials in all LFPs recorded in different sites. Data obtained in two monkeys were pooled at each task epoch (center, movement, hold). Symbols denote data for each recording site. Red crosses represent outlier data. Two-way ANOVA with post hoc Sidak’s test; SR [“epoch” (F_2,99_= 49.085, p < 0.001), “movement direction” (F_1,99_ = 0.355, p = 0.553)], DR [“epoch” (F_2,66_ = 71.78 p < 0.001), “movement direction” (F_1,66_ = 0.02, p = 0.88)]: ***p < 0.001

In total, SR-LFPs were recorded from 27 different sites within the CN (19 sites for Monkey KI and 8 sites for Monkey KE). Of these, most sites (n=19, 71%) produced SR responses without DR responses. This pattern is characteristic of the anatomically defined “cluster area.” In the following, we define the area associated with these sites as the cutaneous area. Only a minority of the sites (n=8, 29%) showed DR responses in addition to SR responses (Figure S3), in line with known properties of the “multimodal” area.^37^

In the cutaneous area, we found that the size of SR-LFPs was suppressed significantly during active movement and during the hold epoch, for both flexion and extension tasks (p<0.001, two-way ANOVA with post hoc Sidak’s test) (Figure 2F). This finding is similar to our previous characterization of the spinal cord,^18^ showing an attenuation of SR-evoked responses during active movement that was however less sensitive to movement direction (flexion versus extension). We also reported similar findings for the intraspinal terminal of cutaneous primary afferents^19, 39^ and for the primary sensory cortex (S1).^18^ Based on those prior findings in combination with our present findings from the CN here, we suggest that movement-related attenuation of cutaneous input is a general phenomenon throughout the dorsal lemniscus pathway (see also^17, 41^ for related findings on primary afferents to the cortex).

In the multimodal area (eight sites), we similarly found that the size of SR-LFPs was suppressed significantly during active movement and during the hold epoch, for both flexion and extension tasks (p<0.001, two-way ANOVA with post hoc Sidak’s test; Figure S4). However, the degree of this suppression was smaller than measured for the cutaneous area in monkey KI (p<0.001, two-way ANOVA with post hoc Sidak’s test; Figure S4). This comparison was only available for monkey KI because we do not have recordings from the multimodal site in monkey KE. The comparison suggests that suppression is a phenomenon predominantly associated with pure cutaneous signals ascending to the medial lemniscus pathway, and that its impact on multimodal signals is more limited.

We recorded DR-LFPs from 12 different sites within the CN. The majority of these sites also produced SR-LFPs (n=8, 66%), with the remaining minority (34%) showing exclusively DR-LFPs. This result confirms a previous report that the representation of muscle afferent inputs is less segregated than that of the cutaneous afferent.^37^ When measuring their amplitude, we also found smaller DR-LFPs during active movement for both flexion and extension (p<0.001, two-way ANOVA with post hoc Sidak’s test; Figure 2 G). Importantly, this finding stands in clear contrast with our previous observations for spinal neurons: in the spinal cord, we reported facilitation of DR responses during extension movements.^20^

Overall, we observed significant sensory attenuation of both cutaneous and muscle-afferent inputs in the CN. The attenuation of sensory signals is scarcely specific for sensory modality and direction of ongoing movement; both SR and DR responses were attenuated, and this attenuation occurred during both extension and flexion movements. At the same time, sensory gain modulation of proprioceptive afferent inputs in the CN is clearly distinct from related effects in the spinal cord; the responses were attenuated in the CN, whereas facilitated in the spinal cord.^20^

### Top-down and bottom-up components of sensory gain modulation

Building upon the findings detailed above, we set out to investigate the source of SR-LFP suppression during movement. Sensory gating is thought to be a process in which bottom-up sensory signals are suppressed by top-down command signals (such as corticospinal motor commands). Since the CN receives not only the bottom-up afference signals, but also top-down corticofugal projections^16, 22–24, 26^, we attempted to dissociate the influence exerted by the top-down commands by comparing the size of SR-evoked LFPs between passive and voluntary hand movements. The rationale behind this approach is that the former configuration (passive) should exclusively reflect bottom-up modulation, while the latter configuration (voluntary) may reflect a mixture of top-down and bottom-up modulations. To achieve passive movement, we applied this manipulation to the hand of monkeys under light sedation, at two different speeds. The manipulandum moved the monkey’s hand from its neutral position to the flexion position along fast (40 degrees/second) and slow (20 degrees/second) trajectories for the two speed conditions. SR-LFP was successfully evoked during different epochs of the resulting movement. Figure S5A shows an example of an SR-evoked LFPs that was significantly suppressed during the movement epoch, with larger suppression for the fast (left) compared with the slow speed condition (right). We repeated the same measurement for LFPs evoked at 22 sites within the CN (18 for monkey KE, 4 for monkey KI). The results again indicated that the size of evoked LFPs was suppressed for both conditions (p<0.001, one sample *t*-test), with the suppression being more pronounced for the faster movement (p=0.004, paired *t*-test; Figure S5B). Because the size of evoked responses was invariant for different static wrist positions (18 sites for monkey KE, p>0.05 on a one-way ANOVA with post hoc Sidak’s test, Figure S5C), the speed-dependent modulation indicates that gating in the CN reflects modulation in signals at the periphery.^22^ Sensory signals from cutaneous and proprioceptive afferents vary in the extent to which peripheral receptors are stimulated,^42–43^ with a dependence on movement kinematics.^44, 52–53^ Therefore, this finding indicates that the ascending cutaneous signals from the hand are passively received in the CN depending on movement kinematics.

Next, we examined top-down effects on sensory attenuation at the level of the CN. To this end, we directly compared the size of SR-evoked responses between active and passive conditions in awake monkeys. In this experiment, we first asked monkeys to perform the task (Figure 2A) while we recorded the precise kinematic performance of the wrist movement as relayed to the manipulandum. We then instructed monkeys to remain still (see Methods), and we manually reproduced comparable movements of the manipulandum. A representative result is shown in Figure 3A and B. We observed suppression of SR-LFPs for both conditions, although the degree of suppression was larger during active movement. This example also shows no clear EMG activity during passive movement. We pooled EMG and wrist angle data across recording sessions (n=13) separately for different movement directions (flexion versus extension), and compared the results between task conditions (active versus passive). EMGs from wrist muscles during wrist movement were significantly larger in the active than in the passive condition (p<0.001, two-way ANOVA with post hoc Sidak’s test, Figure 3C), because there was no clear muscle activity in the passive condition. Agonistic muscles were exclusively activated during active movement (for example, wrist flexor muscles are exclusively activated during wrist flexion). As shown in Figure 3D, movement speed was typically faster in the passive condition (p<0.001, two-way ANOVA with post hoc Sidak’s test, Figure 3D), suggesting that experimenters showed a tendency towards overestimating movement speed when they reproduced the kinematics of active movement using their hand. We combined SR-evoked responses from both flexion and extension trials, and averaged them to produce a plot in the same way as in Figure 2F (Figure 3E). In total (n=13, 9 from Monkey KE and 4 from Monkey KI), we found that suppression was larger in the active condition compared with the passive condition (p<0.001. two-way ANOVA with Sidak’s test). This difference in suppression cannot be ascribed to differences in bottom-up sensory signals between active and passive movement conditions, because movement speed in the active condition was slower, not faster, than in the passive condition (Figure 3D); slower movement speed is expected to produce smaller, not larger, suppression of SR-LFPs (Figure S5B). Based on these results, we conclude that the difference in suppression between active and passive movement predominantly reflects top-down suppression during active movement.

**Figure 3:**
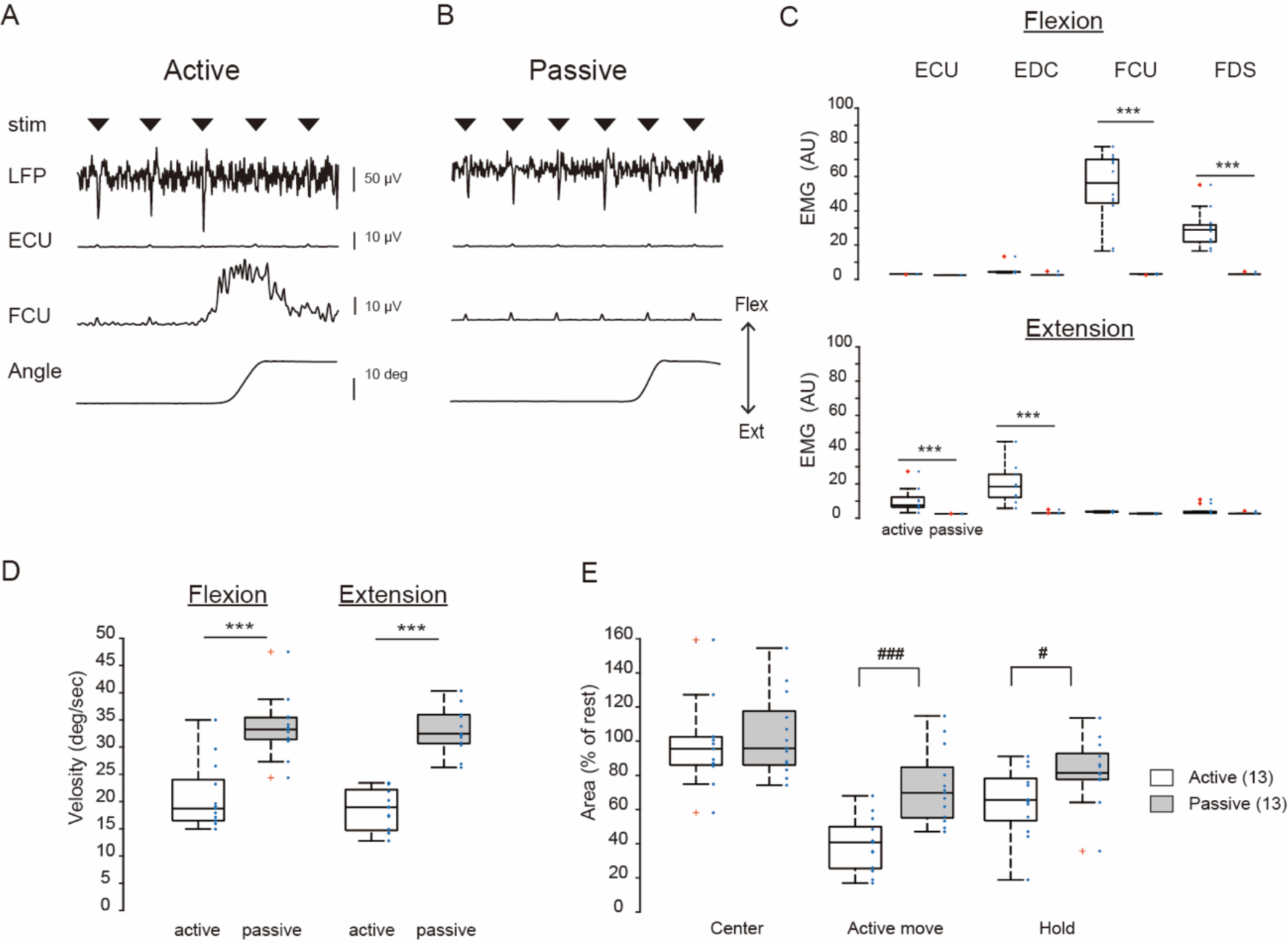
Comparison of SR-evoked local field potentials between active and passive conditions. **A and B**, Typical examples of SR-evoked local field potential (LFP) in the cuneate nucleus during a single active (A) and passive (B) flexion trial. Triangles indicate the timing of the nerve stimuli (stim). From the top, SR-evoked LFP, extensor carpi ulneris (ECU) muscle activity, flexor carpi ulneris (FCU) muscle activity, and wrist angle. Note that there were no muscle activities during passive movement. **C,** The root mean square value of electromyogram (EMG) on ECU, FCU, Extensor digitorum communis (EDC), Flexor digitorum superficialis (FDS) muscle recorded during movement epoch in the active or passive condition. The data was computed at each movement direction and pooled across two monkeys. Symbols denote data for each recording session (same for D and E). Red crosses represent outlier data. In total, we recorded 14 EMGs for FDS, 12 EMGs for FCU, 11 EMGs for ECU, and 10 EMGs for ECU. Two-way ANOVA with post hoc Sidak’s test; FDS [“movement condition” (F_1,44_ = 72.606, p < 0.001), “movement direction” (F_1,44_ =58.24, p < 0.001), “interaction” (F_1,44_ =58.549, p < 0.001)], FCU [“condition” (F_1,44_ = 39.062, p < 0.001), “direction” (F_1,44_ =42.754, p < 0.001), “interaction” (F_1,44_ =39.236, p < 0.001)], EDC [“condition” (F_1,38_ = 27.552, p < 0.001), “direction” (F_1,38_ =17.074, p < 0.001), “interaction” (F_1,38_ =17.074, p < 0.001)], ECU [“condition” (F_1,34_ = 13.842, p < 0.001), “direction” (F_1,34_ =10.632, p = 0.003), “interaction”, (F_1,34_ =10.547, p = 0.003)]:** P = 0.003, *** p <0.001. **D**, Boxplots showing the movement velocity of wrist flexion and extension in active (left) or passive (right) conditions. Two-way ANOVA with post hoc Sidak’s test; [“movement condition” (F_1,48_ = 97.989, p < 0.001), “movement direction” (F_1,48_ = 1.112, p=0.541)]: *** p <0.001. **E,** Boxplots showing the area of SR-evoked field potentials recorded in the active or passive task conditions. Two-way ANOVA with post hoc Sidak’s test; [“movement condition” (F_1,72_ =15.718, p < 0.001), “epoch” (F_2,72_ = 29.472, p < 0.001)]: ### p < 0.001, # p = 0.033

We further confirmed this top-down effect by examining the responses obtained during the pre-movement period (yellow in Figure S6A). We restricted this analysis to trials on which a sufficient number (>12) of stimuli was applied to the SR during this short epoch, to ensure that average SR-LFPs would support adequate signal-to-noise ratios. We analyzed 17 SR-LFPs for the active condition (9 flexions and 8 extensions), and 19 for the passive condition (10 flexions and 9 extensions). We observed SR-LFP suppression during the pre-movement period (p<0.01 on a one sample *t*-test) and during the active movement period (p<0.001) of wrist extension (Figure S6B), however SR-LFPs remained unchanged during the pre-movement period of the passive condition (p>0.05, one sample *t*-test, Figure S6E). These results indicate that top-down suppression of the SR-LFPs already started before the onset of active movement. Although SR-LFPs were typically smaller during the pre-movement period of the flexion movement, this effect was not significant (p=0.077, one sample *t*-test, Figure S6C). We previously documented a similar extension-biased pre-movement suppression of SR-LFPs at the level of the spinal cord in monkeys performing similar wrist movements^18, 39^, suggesting that this extension bias may be specific to the wrist flexion-extension task adopted by those studies. Overall, based on these results, we conclude that CN neurons receive top-down regulation from the motor command that also generates volitional muscle activities.

### Modulation of individual cuneate neurons

We further investigated whether sensory attenuation is also observed at the level of single CN neurons, rather than only at the aggregate activity level reflected by LFPs. It is challenging to achieve selective recording of activity from single neurons in the cluster area of the CN, because many neurons are densely colocalized within this area.^27^ Furthermore, we found that the majority of these neurons produced burst activity with very high firing rates during the awake condition, consistent with a previous report.^45^ Notwithstanding the above difficulties, we successfully recorded activity from 74 cuneate neurons (58 in monkey KI, and 16 in monkey KE). For each neuron, we generated a peristimulus time histogram (PSTH) by aligning the neuronal response to the stimulation pulses applied during the task and assessed the characteristics of the evoked response. Of the 74 recorded neurons, 38 were identified as SR neurons (responded to SR nerve stimulation), 3 were identified as DR neurons (responded to DR nerve stimulation with short latency <10 ms), and the remaining 33 neurons were categorized as not-identified (see methods for details on this classification). We focused our further analysis on the 38 SR neurons.

Figure 4A shows two examples of SR neurons representing different response patterns elicited by SR nerve stimulation: the excitatory (top) and the inhibitory (bottom) patterns, in line with previous findings.^56^ The onset latencies of excitatory and inhibitory responses were 6 ms and 9 ms, respectively. Figure 4B shows the firing rate modulation of two neurons during flexion and extension trials involving voluntary wrist movement. The neuron with an excitatory response to SR stimuli (top in A) showed an increase in firing rate (compared with the center epoch) during flexion (p<0.001, paired *t*-test) and extension movements (p=0.04). The neurons with an inhibitory response (bottom in A) showed decreased firing rate during both flexion (p<0.001, paired *t*-test) and extension (p<0.001). In first case, the firing rate modulation goes in the direction of facilitating cutaneous sensory transmission along the dorsal lemniscus pathway during movement. In the second case, the firing rate modulation goes in the direction of suppressing the transmission. Interestingly, the firing rate suppression associated with the inhibitory response neuron started before movement onset (80 ms before the onset of flexion and 100 ms before extension), suggesting the involvement of top-down modulation. We also found compatible results for a small sample size (n=3) of DR neurons (Figure S7).

**Figure 4:**
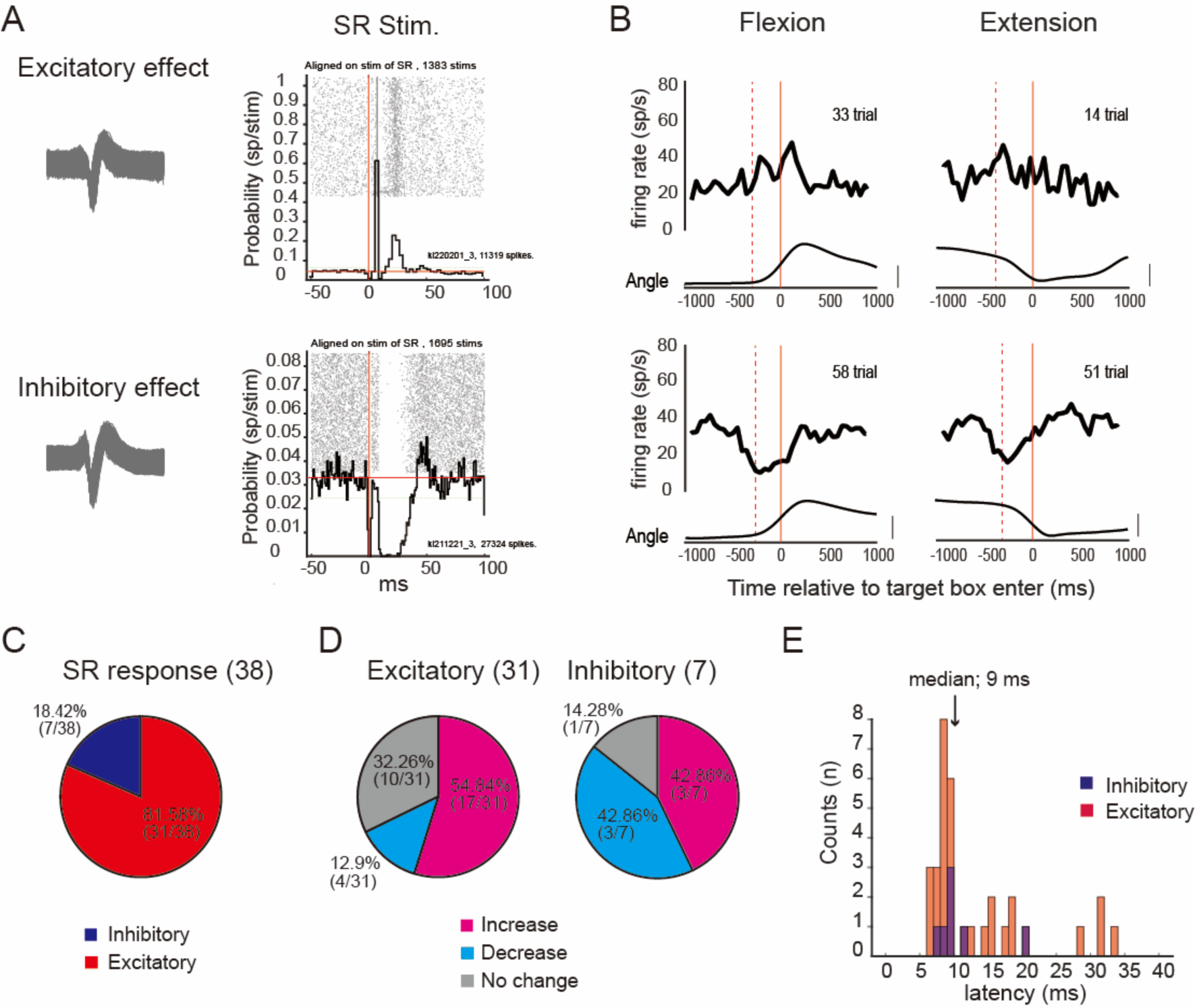
Response patterns of cuneate neurons elicited from SR nerve stimulation. **A,** Examples of the excitatory (**Top**) or inhibitory (**Bottom**) response of the cuneate neuron, elicited by SR nerve stimulation. **Left;** Waveforms of the sample neuron (**Top**: 11319 spikes overlayed, **Bottom**: 27324 spikes overlayed). **Right;** Cell discharge raster plot and its peristimulus time histogram (PSTH) of the sample neuron aligned on the SR stimulation pulses. The vertical red line indicates the timing of the stimulation. In the raster plot, each row represents the order of the response (from older to newer), and each dot represents a timing of action potential detected. The PSTH summarizes the firing profile of the neuron in response to nerve stimulation. PSTH bin size = 1 ms. A solid horizontal red line indicates the mean firing rate during the 100 ms preceding the stimulation pulse. The dashed green line illustrates 2 SDs above or below the mean. **B,** Averaged firing patterns of SR response neurons (the same neurons shown in A) during a wrist flexion/extension task. Excitatory response neurons (**Top**) showed an increase in firing rate during flexion and extension movement, compared with the center epoch (p<0.001 in flexion, p=0.04 in extension, paired t-test). In contrast, neuronal discharge was decreased in inhibitory response neurons (**Bottom**) for both flexion and extension movement, compared with the center epoch (p<0.001 in flexion, p<0.001 in extension, paired t-test). A solid red line illustrates the timing of the entrance of the cursor into the flexion or extension target box. The dashed red line represents the onset timing of wrist movement. **C,** Population of response patterns elicited from SR nerve stimulation. The majority of SR response neurons have excitatory effects (81.58%). **D**, Distribution of discharge patterns of SR response neurons during wrist movement. **Left;** discharge patterns of excitatory effect neurons. **Right;** Discharge patterns of inhibitory-effect neurons. The firing rate of each neuron recorded on the wrist movement phase was compared with that of the rest, and classified as increase, decrease, and no change, respectively (see method). **E,** Onset latency histogram of SR response neurons. Data were from 38 neurons.

Of the 38 SR neurons, 31 (81.6%) showed excitatory responses to SR nerve stimulation, while the remaining 7 neurons (18.4%) showed inhibitory responses (Figure 4C). The majority of neurons, whether associated with excitatory (21/31, 67.7%) or inhibitory responses (6/7, 85.7%), showed significant modulation of firing rate during movement, suggesting that these CN neurons play a significant role in actively modulating the transmission of peripheral sensory signals along the dorsal lemniscus pathway. The onset latency of these SR neurons ranged between 6 ms and 34 ms, with 17 neurons (44.73%) showing a response latency below the median value of 9 ms (Figure 4E). Among the latter neurons, those with excitatory responses exhibited a sharp and narrow peak with few jitters (as shown in Figure 4A), which is a characteristic feature of the monosynaptic projection from primary afferents^59^. It is therefore likely that these short latency neurons received direct, monosynaptic input from SR afferents, although some CN neurons may be activated through indirect pathways, such as the postsynaptic dorsal column pathway.^21^

Next, we examined the modulation of response probability in the SR neurons with excitatory responses (31 neurons identified in Figure 4C) while stimulating the SR nerve during both active and passive movements (Figure 3). Of the 31 SR neurons, 7 were excluded from subsequent analysis because of insufficient numbers of stimuli or responses to support adequate estimation of response probability for each movement epoch (see Methods). We calculated response probabilities for the remaining 24 SR neurons, of which 15 were recorded during active movement, 5 were recorded during passive movement, and 4 neurons were recorded during both active and passive movements. We counted these 4 neurons twice when assigning their data to active and passive conditions. Following this procedure, the active condition counted 19 neurons, and the passive condition counted 9 neurons.

Figure 5A–C shows three representative neurons showing modulation of their evoked response elicited by SR nerve stimulation, under active (A and B) and passive (C) movement conditions. In these neurons, we observed a distinct response during the center epoch (D and F, shorter latency; E, longer latency), however their response decreased during both active and passive movements (G–I). Background activity for these neurons remained unchanged (p=0.0543 in D, p=0.0973 in E, paired *t*-test) or increased (p=0.009 in F) during the movement epoch compared with the center epoch, suggesting that these suppressive effects could not be explained by a general decrease in excitability. Rather, we suggest that an active inhibitory mechanism, which is selective for the transmission of signals from cutaneous afferents to these neurons, underlies the movement-related attenuation of the SR response.

**Figure 5:**
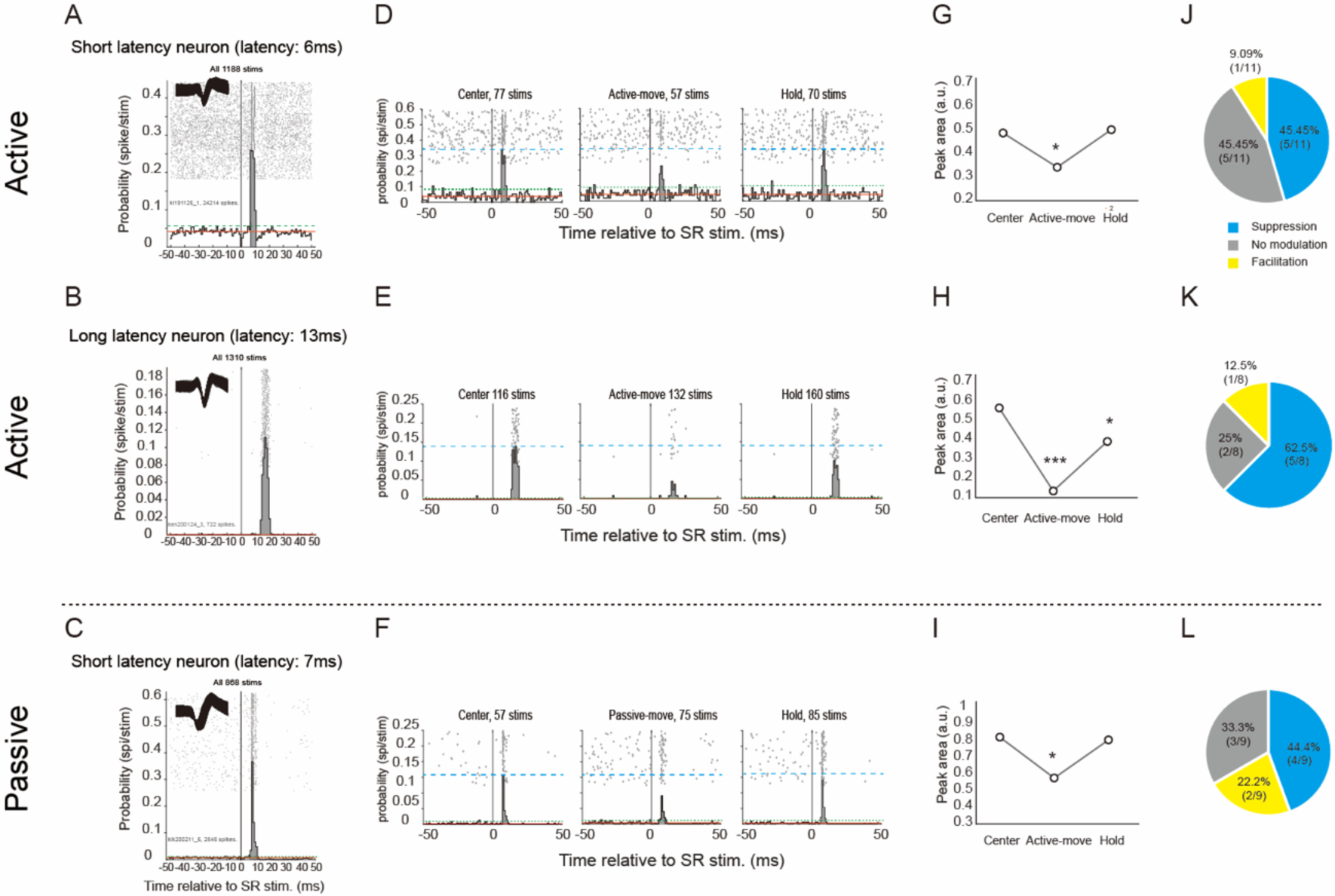
Modulation of SR-evoked response in the cuneate neuron during a motor task. **A-C,** Raster plot and peristimulus time histogram (PSTH) of three cuneate neurons aligned on all SR nerve stimulation pulses during the active condition in a short-latency neuron (A), in a long latency neuron (B), and during the passive condition (C). All formats are similar to Figure 4A, but the measured peak area was filled with gray. The peak area was defined as where it crosses 2 SDs (dashed green line) above the average of the baseline. Insets indicate the waveform of a sampled neuron. **D-F,** Raster plot and PSTH of the sample neurons shown in A-C were compiled separately for three task epochs. From left, “Center”, “Movement”, and “Hold” epoch. Numbers in each graph indicate the number of stimulation pulses. The Blue dashed line indicates the peak value of evoked responses in the center period. **G-I,** Difference in peak areas among three behavioral epochs of three sample neurons. Note that SR-evoked responses are clearly suppressed during the movement period in either behavioral condition. * p<0.05, *** p<0.001 significant difference in peak area between the center and another epoch by Binomial test. **J-L,** Summary of modulation patterns in the excitatory response neurons. About half of the response neurons, in both short- and long-latency neurons, showed suppression in the SR evoked responses during movement epoch in both movement conditions.

We repeated this analysis for all excitatory response neurons (n=24). We present results separately for the modulation observed during active movement in neurons with short (<9 ms) latency response (Figure 5J, n=11), those with longer latency (>=9 ms, Figure 5K, n=8), and for the modulation during passive movement (Figure 5L, n=9). During passive movement, 5 neurons responded with short latency, and 4 neurons with longer latency. We found that the majority of neurons in all groups showed a reduction of the peak area associated with SR-evoked responses during both active and passive movements (blue in Figure 5J–L). Furthermore, the proportion of neurons with decreased response probability to SR stimuli is higher in the active condition (10/19, 52.63%) than in the passive condition (4/9. 44.4%). These results indicate that the cutaneous input to cuneate neurons is modulated during both active and passive movements, and that the degree of suppression is greater in active movement than passive movement, confirming our findings with SR-LFPs (Figure 3).

Interestingly, none of the neurons with decreased response probability to SR stimuli (n=10 in the active condition and n=4 in the passive condition) showed a decrease in firing rate during movement: during the active condition, 4/10 neurons increased their firing rate and 6/10 neurons showed no change, while in the passive condition 3/4 neurons increased their firing rate and 1/4 neurons showed no change. Because we restricted our analysis to neurons that showed similar mean firing rates throughout the task epoch or an elevated mean firing rate during the movement epoch (see Methods), this result implies that the selective suppression of cutaneous inputs is independent of the general excitability of the receiving neurons, as we also found in representative data (Figure 5A–C).

We previously reported a similar finding for SR inputs to spinal first-order interneurons: short-latency inputs to these neurons were suppressed, while the overall neuronal excitability increased.^19–20^ Our subsequent work indicated that the suppression of response probability was partly mediated by presynaptic inhibition of the primary afferent terminal from descending motor commands.^19^ We therefore suggest that presynaptic inhibition may mediate part of the response suppression found in cuneate neurons, especially for early onset neurons (Figure 5A).

## Discussion

A hallmark of hand movement is the rich somatosensory feedback associated with this action. To fully understand the neural control of movements, it is critical that we understand how this feedback information is processed by the brain. The CN represents one of the first entry points for hand somatosensory afferents to the CNS. For this reason, the modulation of sensory signals at the level of the CN has an impact on sensory processing by subsequent neural structures. However, our current knowledge about sensory processing in the somatosensory network is restricted to higher brain areas,^46–47^ with the potential contribution of the CN remaining largely unknown. In this study, we found that both cutaneous and proprioceptive signals were suppressed in the CN during voluntary movements. Because this suppression was observed within a region of the CN where afferent signals are sent to cuneothalamic relay neurons,^23, 40, 48^ it is likely that S1 and higher cortical structures operate on somatosensory signals that have already been pre-processed by the CN, rather than on genuine afferent signals.

### Nonspecific suppression of somatosensory signals within the cuneate nucleus

We found that sensory attenuation at the CN was not specific for input modality (cutaneous versus muscle afferents), nor for movement direction (flexion versus extension; see Figure 2). This finding suggests that the CN plays a role in suppressing some afferent signals irrespective of movement or sensory modality. Similar to the over-trained wrist movement adopted in this study, the performance of proficient movements may rely primarily on sensory prediction signals generated by the internal model,^11^ rather than on actual afferent signals. Under this scenario, sensory attenuation at the level of the CN plays a central role in regulating unnecessary afferent signals generated by one’s own movements, and facilitates efficient sensory computation by higher centers.

This novel finding stands in clear contrast with our previous characterization of the spinal cord. In this structure, DR inputs to first-order interneurons are enhanced during wrist extension movements and unaffected during flexion movement,^20^ while SR inputs are consistently suppressed during both movements.^18–19^ Therefore, sensory attenuation at the level of the spinal cord is more selective than its counterpart in the CN. This difference may originate from known functional differences between the two structures: the CN is only involved in the transmission of somatosensory signals to higher centers, while the spinal cord is also involved in turning these signals into motor outputs via reflex pathways. Here, we suggest that the attenuation of redundant afferent signals during voluntary movement may represent a general mechanism that is implemented at various sensory relay stations throughout the CNS, as we found it at the level of both spinal cord and CN. Importantly, however, we also found facilitation of SR-evoked responses during movement for a small fraction of the neuronal population (Figure 5J; see also related results by Versteeg and collaborators^49^). Therefore, CN neurons also amplify task-relevant afferent signals as input to reflex-related neurons in the spinal cord. In this sense, the CN probably functions as a flexible gain controller of somatosensory afferent signals.

### Gating of afferent signals at the level of spinal cord, cuneate, and somatosensory cortex

In the dorsal column lemniscus pathway, afferent signals reach S1 via tri-synaptic or quadri-synaptic linkages. They are relayed by the dorsal horn in the spinal cord, the cluster area in the CN, and the VPL nucleus in the thalamus. In the past, we have compared the degree of suppression for SR-evoked responses between the spinal cord and the S1 in monkeys performing a similar wrist flexion-extension task. We found that suppression was more pronounced in the S1 (60% reduction in the magnitude of SR-LFPs compared with the control condition) than in the spinal cord^18^ (20% reduction). The present study measured comparable reductions for the CN of about 59.79% and 60.69% (Figures 2 and 3). The degree of suppression in the CN was therefore similar to that observed in S1, but larger than measured in the spinal cord. The latter difference may reflect three distinct mechanisms. First, as already discussed, the reduced sensory attenuation of the spinal cord may be explained by its additional role in modulating reflex outputs that may not be coherent with the modulation carried by the ascending pathway. Second, the suppressed afferent signals transmitted by spino-cuneate neurons were further suppressed in the CN (summation effect). Third, the extent of suppression was indeed larger in the CN. This result, combined with the comparable reduction of SR-LFPs observed for CN and S1, indicates that further attenuation by subsequent relay stations at thalamic and S1 levels may be relatively subtle. This interpretation suggests that the sensory attenuation reported for cortex simply reflects attenuation at the level of CN and spinal cord.

Interestingly, when we restrict our focus to the cutaneous area (putative cluster area), the primary area that relays cutaneous signals from CN to thalamus, we found a large reduction of SR-LFP amplitude at more than 60% compared with the control condition (71.6% reduction from the rest condition, Figure S4), which was larger than observed in S1 (60%). These results support the notion that somatosensory signals may be amplified after passing through CN.^17^ It is therefore possible that the redundant component of the afferent signal is first suppressed by the filter mechanism implemented within spinal cord and CN, and that the residual, meaningful sensory signal is then amplified by higher sensory relay stations for improving somatosensory acuity^50^ during voluntary movement. Further studies that simultaneously compare the strength of sensory gating at each relay station under the same behavioral task would be required to confirm this hypothesis.

### Bottom-up and top-down modulation of somatosensory signals

We showed that both top-down and bottom-up signals (Figures 3, S5, and S6) play a role in mediating sensory attenuation in the CN of monkeys performing voluntary movement. This conclusion is in good agreement with previous work suggesting that bottom-up inputs act as a filter of afferent signals and top-down inputs act as sources of sensory gating in the CN.^16–17, 22–24, 26, 29–30^ An outstanding question is their relative role in behavioral control. We suggest that bottom-up attenuation facilitates the extraction of dynamic information from the moving body, while top-down attenuation aids the extraction of unpredicted afferent signals, and that both tasks are achieved via attenuation of redundant afferent signals.

When we move our hand, a specific skin region is stimulated, and a substantial number of mechanoreceptors is activated.^1, 51–52^ A ubiquitous proprioceptor, i.e., Muscle spindle and Golgi tendon, in the hand is also activated at the same time,^53–54^ leading to a large influx of somatosensory signals to the CN. Without proper regulation of this afferent bombardment, the sensory system may be overwhelmed by excessive information. As a consequence, it may not accomplish successful recognition of limb dynamics, which is crucial for accurate movement control. It is therefore reasonable to expect that attenuation of incoming signals would happen early along the somatosensory hierarchy.^55^ The CN is a good candidate for taking on this early regulatory role, because it supports convergence of multiple peripheral inputs.^56^ The extraction of haptic features achieved within the CN by integrating signals from mechanoreceptors^57, 58^ supports this hypothesis.

As already discussed, afferent signals produced during voluntary movement can be predicted within the internal model^11^ when performing skilled movements. Once afferent signals representing active body dynamics have been extracted via bottom-up attenuation, they could be further attenuated by top-down mechanisms in the CN for prioritizing sensory prediction over noisy afferent signals.

### Presynaptic and postsynaptic inhibition of cuneate neurons

We showed that the neuronal activity of SR neurons evoked by peripheral stimulation was inhibited during the movement phase, even though their task-related firing rate was increased or unchanged. In the spinal cord, we initially reported a similar observation: the monosynaptic input from sensory afferents to postsynaptic cells is decreased, while the total output of postsynaptic cells is facilitated or unchanged during voluntary movement.^19^ We subsequently discovered that suppression is partly induced by presynaptic inhibition on afferent terminals from descending motor commands.^39^ In the case of the CN, it is currently difficult to conclude, in a classical sense, that the early onset CN neurons (Figure 5A) receive monosynaptic connectivity from SR afferents: we were unable to measure the “segmental” latency of individual responses because of difficulties associated with recording incoming volleys within the CN. However, we measured higher response probability with minimal jittering in response delay (Figure 5A), which strongly suggests that these responses represent the monosynaptic connection.^59^ We then suggest that the suppression in responsiveness to SR inputs for these neurons (Figure 5A) may be, at least in part, generated by presynaptic inhibition of afferent terminals within the CN, as reported in other species.^22, 25, 30, 60–62^. With regard to the sensory attenuation found for late-onset neurons (Figure 5E, middle), suppression through local postsynaptic inhibitory interneurons^22, 29^ may represent a more plausible pathway for mediating this mechanism.

## Methods

### Subjects

We obtained data from two male Macaque monkeys (monkey KE: Macaca fuscata, 9.5 kg, aged 10 years at the time of recordings; monkey KI; Macaca mulatta, 8.5 kg, aged 9 years). All experimental procedures were approved by the Institutional Animal Care and Use Committee at the National Institute of Neuroscience, National Center of Neurology and Psychiatry, Japan. During training and recording sessions, the monkeys sat upright in a primate chair with the right arm restrained and the elbow bent at 90°. The monkey’s right hand was held in a cast, with fingers extended and the wrist in the mid-supination/pronation position. The cast was attached to a servomotor-driven manipulandum (Techno-hands Co., Ltd, Yokohama, Kanagawa) that measured flexion-extension torque and angle about the wrist. The left arm was restrained loosely to the chair.

### Task paradigm

Each monkey was trained to flex or extend his right wrist in accordance with an instruction, as described below (wrist flexion-extension task; Figure 2A). A wrist flexion-extension torque applied to the manipulandum controlled the position of a cursor displayed on a video monitor in front of the monkey, using TEMPO system (Version 11.7, Reflective Computing, WA, USA). Because the monkeys performed the task with their right hand, a wrist flexion led to a leftward displacement of the cursor. Trials began with the monkey holding the cursor in a center target window corresponding to zero torque for 0.8–1.2 s (Center hold). Next, flexion or extension targets (empty rectangles) were shown to the left or right of the center target, indicating the sign of movement start (GO signal). Following the GO signal, the monkey moved the cursor to the desired target (Active movement) and held the cursor in the target window for a period of 0.8–1.0 s (Active hold). Movements were performed against an elastic load applied by the servomotor (10 N×m). The monkeys moved their hand over a range of 10 to 20 degrees. At the end of the active hold period, the peripheral target disappeared. The monkey then relaxed its forearm muscles, allowing the servo-spring to passively return the wrist to the zero-torque position (Passive back). At this point in time, the monkey was rewarded with apple sauce for successful trials (Reward). At the end of the reward time window (0.8–1.0 s), the next trial started after a 0.8–1.0 s interval (Rest). When the monkey failed Center hold, Active movement, or Active hold during any of the above steps, the trial was aborted and marked as an error trial. On each trial, Center hold and Active hold times were changed at random within a 200-ms window to avoid moving the hand predictably. In a separate session, we also trained monkeys not to resist while the manipulandum reproduced a movement comparable with the active movement (passive condition). The experimenter was tasked with producing a movement trajectory that mimicked as closely as possible the trajectory enacted during active movement, by manually moving the manipulandum (Figure 3). The four epochs marked in gray in Figure 2A were used for analysis (Rest, Center Hold, Wrist movement, Hold). In addition to these task sessions, we included an additional passive movement session under light anesthesia with xylazine hydrochloride (0.6 mg/kg) and butorphanol tartrate (0.05mg/kg). During this session, the monkey hand was automatically flexed by a manipulandum with fast (40 degree/second) and slow (20 degree/second) speed conditions over a range of 20 degrees. To confirm the influence of wrist position, we recorded neural responses by keeping the monkey’s hand at 20 degrees of flexion, neutral, and 20 degrees of extension.

### Surgical implant

After the monkeys had learned the required task, surgeries were performed aseptically with the animals under 2.0%–3.0% sevoflurane anesthesia with a 2:1 ratio of O_2_:N_2_O. Pairs of stainless steel wires (AS631, Cooner Wire) were implanted subcutaneously in six muscles: extensor carpi ulnaris (ECU), extensor carpi radialis (ECR), extensor digitorum communis (EDC), flexor carpi radialis (FDS), flexor carpi ulnaris (FCU), and flexor digitorum superficialis (FDS). Each muscle was identified based on its anatomical location and characteristic movements, elicited by trains of low-intensity electrical stimulation applied directly on the muscle fascia (0.5-ms bipolar pulses, 50–100 Hz, 20–30 trains). These muscles were active in either flexion or extension direction when monkeys performed the task (Figure 3C). A nerve cuff electrode (tripolar) was implanted on the superficial branch (SR) and the deep branch (DR) of the radial nerve for stimulating cutaneous and muscle afferents (Figure 1C): SR exclusively innervates a patch of skin on the dorsal, radial aspect of the hand, while DR mostly innervates muscle fibers and afferent fibers of extensor muscles and its tendon organ in the forearm. Another nerve cuff electrode (tripolar) was implanted on the radial nerve proximally, ∼10.0 cm apart from the SR and ∼2.0 cm from the DR, for recording conducting volleys produced by SR or DR stimulation (Figure S2C and D). EMG and nerve cuff connectors were cemented to the skull with dental acrylic that anchored to the bone via screws. A custom-made recording chamber (polyetherimide, a circular cylinder with a diameter of 40 mm) was implanted over a craniotomy in the left side of the skull.

### Recording procedure

During recording sessions, we restricted head movement by securing the monkey’s head with a thermoplastic mask (Uni-frame, Toyo Medic, Tokyo, Japan) adjusted to fit each monkey.^63^ An hydraulic micromanipulator (MO-973; NARISHIGE, Japan) was attached to the recording chamber via a custom-made X-Y positioning stage. Neuronal activity in the CN was recorded extracellularly with glass-insulated tungsten single microelectrodes (Alpha Omega Engineering Ltd., Israel) or platinum/iridium liner microelectrode arrays (8 or 16 channel S-probe; inter-electrode spacing of 100 or 200 µm; Plexon Inc., USA) while the monkey performed wrist flexion and extension tasks. The range of impedance for a single-channel electrode was between 0.5 and 1.0 MΩ, and for a S-probe between 0.3 and 1.0 MΩ at 1 kHz. We adopted the following procedure for recording from the CN. First, we calculated a distance from the surface of the cortex to the medulla oblongata based on MRI and CT images (Figure S1A). Next, we moved an electrode over the calculated distance while monitoring neural activity on the way to the cuneate nucleus. Once neuronal activity went silent as we reached the fourth ventricle, this event was used as a landmark for estimating the location of the electrode tip. We then carefully moved the electrode while monitoring the neural response to SR or DR stimulation. When we found a receptive field receiving input from the forearm area, we waited at least 5 minutes for unit activity to stabilize and then started recording. We aligned the recording sites within the CN based on the depth of the electrode tip at which unit activity was first recorded during each penetration (Figure S1C). The SR or DR was stimulated (bipolar pulses of 0.1 ms) at a constant frequency of 2 Hz during recording sessions. The threshold current that evoked an incoming volley from SR or DR stimulation was recorded during each session. Stimulus intensity was set at 1.5–2.0 times the threshold for SR stimulation, and 1.1–1.2 times the threshold for DR stimulation. We ensured the incoming volley evoked by a given stimulus intensity was stable for the time during which we obtained the data (Figure S2C–D).

Data were amplified (×20) and digitized by AlphaLab SNR (Alpha Omega Engineering Ltd., Israel) at different sampling rates. The rates for sampling neuronal activity were 22 or 44 kHz, those for EMGs were 22 kHz, and those for wrist torque and angle were 2.75 kHz. Output from the amplifier was bandpass filtered for online monitoring of LFPs (5–200 Hz) and action potentials (200–9000 Hz) from the CN. An Ag-AgCl ball electrode placed on the scar tissue overlying the cortical surface.

For SR-or DR-evoked LFPs, signals were bandpass filtered (2^nd^-order Butterworth filter between 5 and 100 Hz) and down-sampled to 1100 Hz. For single- or multi-unit activities, signals were bandpass filtered between 200 and 9000 Hz. Action potentials were isolated from background activity online using a template-matching spike-shorting algorism (AlphaLab SNR). Because it is difficult to ensure the quality of isolation for multichannel electrodes, we monitored unit activity from a specific channel to ensure recording stability. Spiking activities were manually sorted offline using spike sorting software (Offline Sorter 4.11, Plexon Inc, USA) to improve the isolation of each unit. Offline isolation quality was assessed by compiling an inter-spike-interval (ISI) histogram. The stationarity of single cell activity was checked by inspecting the nerve stimulation response and task-related firing modulation of each cell. Cells that showed pronounced trial-by-trial variance for these parameters were omitted from the study.

For EMGs, signals were bandpass filtered (2^nd^-order, Butterworth filter, between 50 and 2500 Hz), rectified, lowpass filtered (cutoff of 20 Hz), and then down-sampled to 100 Hz. The angular position of the wrist joint was recorded along with other signals and down-sampled to 100 Hz.

### Data analysis

We assessed the epoch-dependent modulation of sensory-evoked responses and the task-related neuronal activity in the CN during wrist flexion and extension tasks. At the beginning of the assessment, we redefined each task epoch by analyzing EMG activities and the movement trajectory of the wrist joint. First, the EMGs for each muscle and movement trajectory were aligned to the entrance timing of movement cursor into flexion or extension target box one second before and after, and then averaged within the session. We calculated the mean of EMG amplitude and angle of the wrist joint between 1000 and 500 ms before the entry time point of the target box, while visually verifying that there was no EMG activity within this period, which was used as baseline of the pre-movement period. The onset of wrist movement and EMG activity was defined as the time point at which the relevant characteristic crossed four standard deviations (SD) of the mean of the baseline. For each experimental session, we computed the onset of EMGs for each muscle, and selected the earliest one from a set of muscles that activated during wrist movement. Movement velocity was computed from the displacement of wrist angle over 100-ms epochs before and after entering the target box, in each task condition (Figure 3D).

After redefining the onset point of wrist movement and EMG activity, we defined four movement-related epochs: (1) Rest, the 800-ms interval before center box turns on; (2) Center, from the onset of the center target to 100 ms before the flexion/extension target box turns on; (3) Movement, from onset of wrist movement to 100 ms after target box enters; (4) Hold, from 100 ms after the entry into the target box to the time point at which the flexion/extension target box turns off. In addition to these epochs, we also defined (5) Pre-movement epoch, from 100 ms before EMG onset to movement onset, and (6) Late center, the 300-ms interval before the initial point of pre-movement. We analyzed only trials in which the monkey was rewarded, and i.e., successful trials, and trials in which all behavioral events were adequately recorded.

The SR- or DR-evoked LFPs were compiled and averaged separately for each behavioral period to study the associated modulations depending on behavioral context. We measured onset latencies and size of evoked LFPs as illustrated in Figure S2A–B. Briefly, for SR stimulation (Figure S2A) we measured onset latency of the evoked LFP from stimulus onset to the onset of the downward deflection of the earliest component of the LFP. We measured the size of evoked potentials using the peak area under the baseline from the onset to the offset of the averaged waveform. The peak onset was defined as the time at which the waveform following the stimulation pulse crossed 2 SDs below the average of the baseline. The peak offset was similarly defined as the time at which the waveform crossed the above threshold in the opposite direction.

Baseline activity was measured from 50 to 20 ms before stimulus delivery. For DR stimulation (Figure S2B), onset latency of evoked LFPs was measured using the same procedure adopted for SR stimulation, however size was measured using a different procedure. To avoid contamination of reafference signals coming from muscle contraction caused by DR stimulation, the size of the DR-evoked LFPs was measured using the peak area under the baseline from the onset to the peak of the averaged waveform. Figure S1B shows electrode insertion points over the cortical surface within the recording chamber. For both SR and DR stimulation, the minimum number of stimulation pulses was arbitrarily set at 12 stimulations per epoch, to obtain reliably evoked LFPs for each epoch.

We generated peristimulus time histograms (PSTHs) from individual neurons aligned to the timing of the stimulation pulse. The PSTHs included data from 50 ms before to 100 ms after the stimulation pulse, with a bin size of 1 ms. The method we used to compute the peak amplitude was similar to that used by Seki et al.^19^ First, we computed the mean evoked response by pooling all stimulation pulses recorded during the task so as to maximize the signal-to-noise ratio. Next, the baseline firing rate was computed as the mean bin height within the 100-ms epoch preceding the stimulation pulse. We then used onset and offset points of the mean evoked response to compute the peak area evoked during each task epoch. The peak onset was defined as the time at which the PSTH following the stimulation pulse crossed 2 SDs above or below the average of the baseline firing rate. Similarly, the peak offset was defined as the time at which the PSTH crossed 2 SDs around the average a second time. For SR stimulation, when the recorded neuron presented a positive peak area above 2 SDs from the baseline, it was classified as an excitatory response neuron. Similarly, if the neuron presented a negative peak area below 2 SDs from the baseline, it was classified as an inhibitory response neuron. For DR stimulation, we defined recorded neurons as DR neurons if they responded to DR stimulation with a short latency of less than 10 ms. This criterion is designed to eliminate contamination by reafference signals coming from muscle contraction caused by DR stimulation. The peak area was computed as the sum of the bins above or below the baseline firing rate during the peak duration. Thus, the peak area can be understood as the mean number of spikes above the baseline evoked by each stimulation pulse during the peak time window.

To characterize the discharge patterns of cuneate neurons during wrist movement, we computed discharge histograms with a bin width of 20 ms and aligned to the entry time of flexion or extension target box, which ranged between 1000 ms before and 1000 ms after entry of the target box (Figures 4B and S7). The mean firing rate of each task epoch (rest, center, movement, and hold) was calculated to determine whether recorded neurons were responsive or silent. The discharge patterns of cuneate neurons were categorized as a combination of facilitation (increase) or suppression (decrease) when their firing rate during the Movement or Hold epoch was significantly increased or decreased compared with that of the center. If there was no significant change in firing rate between Center and other epochs, the neurons were categorized as presenting no modulation (no change).

To assess the modulation of responses from cuneate neurons, we evaluated the epoch-dependent modulation of peak area and firing rate during the wrist flexion-extension task. We restricted the analysis to behavioral epochs during which a sufficient number of stimulus pulses was applied, to obtain an unbiased estimate of the evoked response area. The minimum number of stimulation pulses was arbitrarily set at 12 stimulations per epoch, as a compromise between the reliability of the peak area and the size of the database.

Although the aim of this study was to examine how evoked responses are influenced by behavioral parameters like task epoch or movement direction, the response could also be affected by the overall excitability (firing rate) of a given neuron. Because the firing rate for each neuron was typically different between epochs, the impact of these differences on our interpretation of the associated response modulation is not negligible. For these reasons, when studying the response probability of cuneate neurons during SR stimulation, we restricted our analysis to neurons that maintained a relatively stable baseline firing rate throughout the task epoch, or an elevated baseline firing rate during the movement epoch. This was achieved by comparing the mean baseline firing rate between epochs. We then evaluated the differences in evoked response area across task epochs.

### Statistics

For each task session, the area of SR- or DR-evoked LFPs was calculated based on movement direction and compared between rest and other behavioral epochs (late center, pre-movement, movement, or hold) using a one-sample *t*-test. We analyzed group data for the area of evoked LFPs during SR and DR stimulation across three behavioral epochs (center, movement, and hold) using a two-way analysis of variance (ANOVA) test with factors “epoch” and “movement direction.” To test the significance of size effects associated with evoked LFPs during SR stimulation between active and passive conditions, we used a two-way ANOVA test with factors “epochs” and “movement condition.” EMGs and movement velocity between active and passive conditions were also compared using two-way ANOVA tests with factors “movement condition” and “movement direction”. The amplitude of incoming afferent volleys evoked by SR or DR stimulation was analyzed by one-way ANOVA tests. To compare SR-evoked LFPs between recording areas (cutaneous versus multi-modal), we used two-way ANOVA tests with factors “area” and “epoch.” The size of SR-evoked LFPs obtained under light sedation was compared between center and movement epochs using one-sample *t*-tests. We used paired *t*-tests to assess the effect of movement speed on the size of evoked LFPs during SR stimulation. We analyzed the size of SR-evoked responses for each wrist position using one-way ANOVA tests. If a significant effect was detected, we used the Sidak’s post hoc test to correct for multiple comparisons. The size of the peak area in single unit responses was analyzed by binomial test. The mean firing rate of cuneate neurons during each task epoch and the baseline firing rate from PSTHs were compared between center and other behavioral epochs (movement and hold) using paired t-tests. Except when stated otherwise, measurements are presented as mean ± standard deviation. P values <0.05 were considered significant in all statistical analyses. We used the Matlab_R2019b statistical toolbox (MathWorks Inc., Natick, USA) to perform binomial tests and paired *t*-tests for the analysis of firing rate modulation. We used SPSS version 22 software (IBM SPSS, IBM Japan Ltd, Tokyo, Japan) to perform the other tests.

## Acknowledgements

This work was supported by Grants-in-Aid from the Japan Society for the Promotion of Science (JSPS), grant numbers 19H05724, 19H01092, 26120003, and 23H05488 to K.S., and grant numbers 17J05310, 19K19898, 22H03500, and 23H04372 to S.K. S.K. was supported by a Research Fellowship from the JSPS.

We thank K. Oida for animal care and technical assistance, and M. Kudo for specimen preparation.

## Author contributions

K.S. planned the experiments, S.I. and H.G. developed task devise, S.K., C.S., S.K., and T.O. performed the physiological experiments, S.K. analyzed the data, and S.K. and K.S. wrote the paper. All authors contributed to generating the draft and final version of this paper.

---

**Figure S1:**
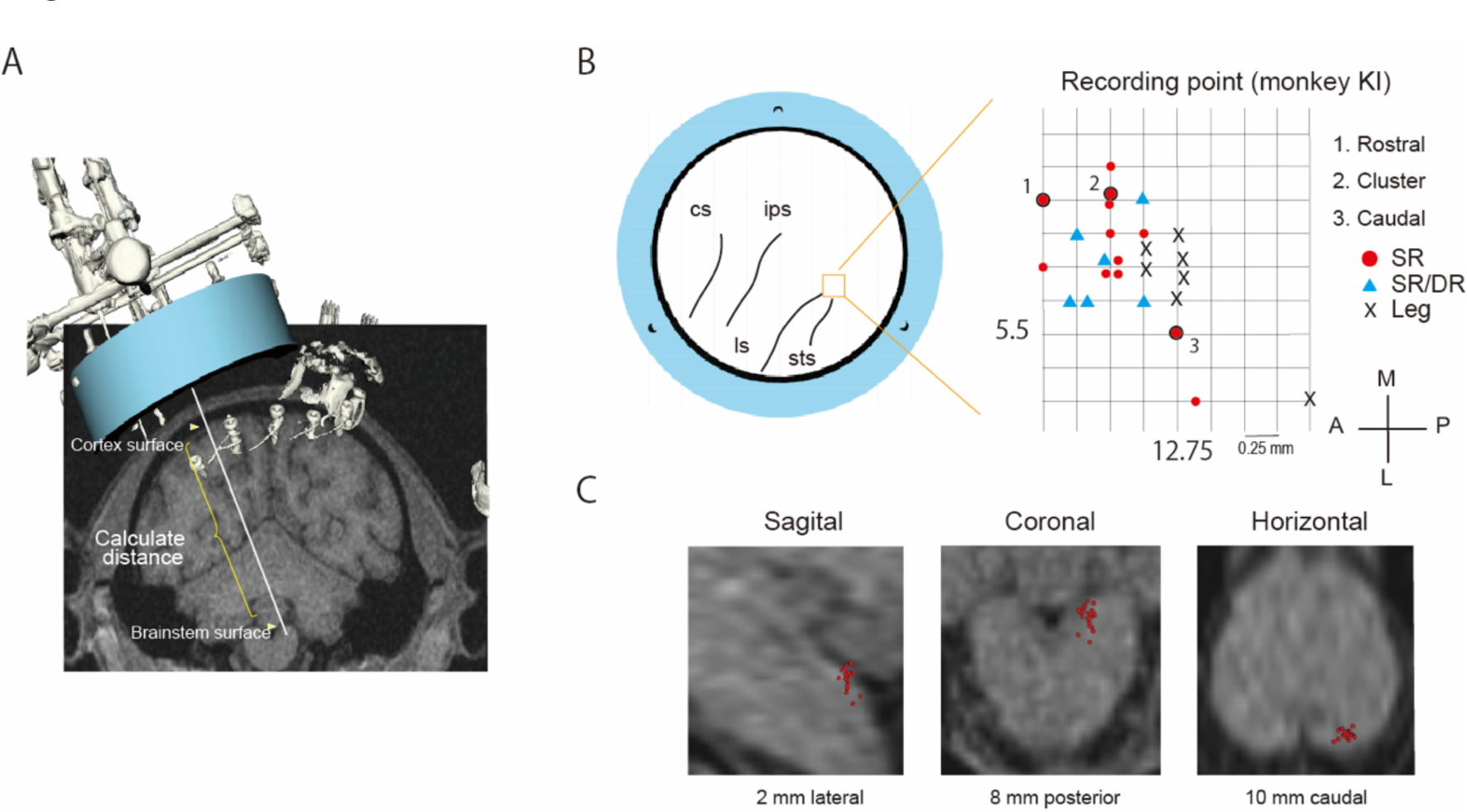
Recording method. **A,** Coronal view of MRI image aligned with recording chamber and recording electrode reconstructed from CT data. A distance from the surface of the cortex to the medulla oblongata (marked by a white triangle) was calculated using Cicerone software^38^. **B,** Electrode insertion point map on the recording chamber in monkey KI. The recording chamber was tilted 25 degrees from the horizontal plane. Red circles indicate the point in which only SR-evoked response was recorded. Blue triangles indicate the point in which not only SR-evoked response but also DR-evoked response was recorded (multimodal area). Cross marks indicate the point at which the responses to a light touch of the leg were observed. The number 1 to 3 represent the respective points shown in Fig.1 D-E (1: Fig.1D 0.5 mm rostral point, 2: Fig.1E Core, 3, Fig.1F 0.5 mm caudal point). A: anterior, L: lateral, M: medial, P: posterior. **C,** Enlarged view of MRI image with the estimated recording point in the cuneate nucleus. Red dots represent each recording points which overlaid a representative MRI image. The sagittal view was 2 mm lateral from the midline. The coronal view was 8 mm posterior from the interaural line. The horizontal view was 10 mm caudal from the interaural line.

**Figure S2:**
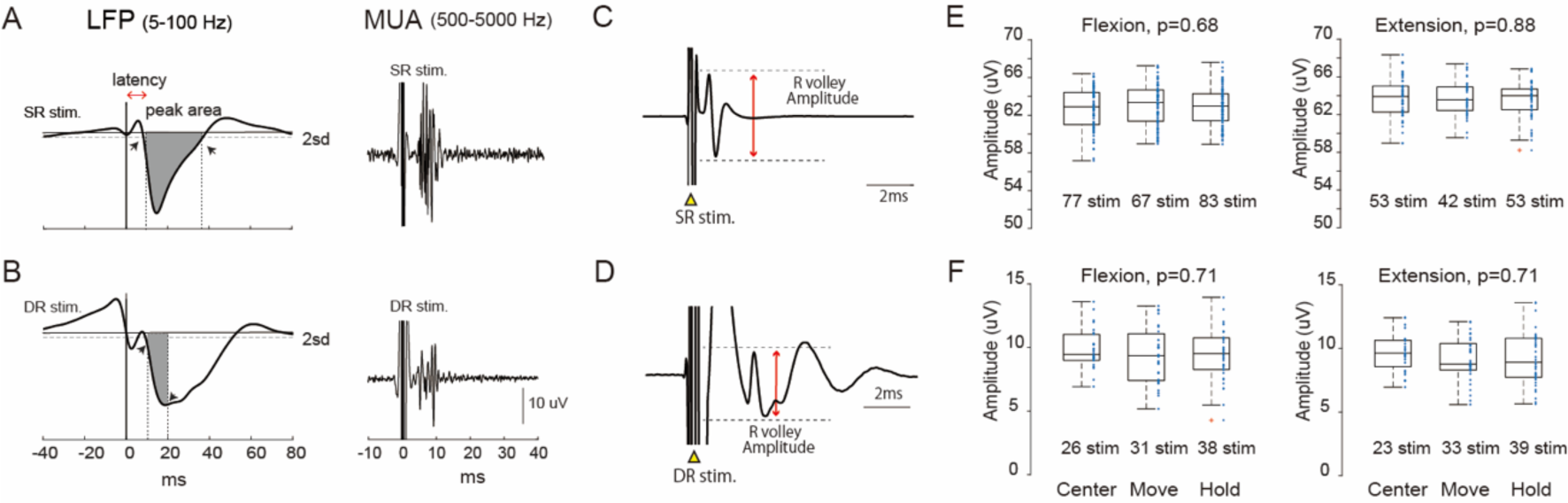
The measurement of onset latency and size of evoked potentials. **A,** An example of SR-evoked local field potential (LFP, **Left**) and multiunit activity (MUA, **Right)**. The onset latency of evoked potentials (red arrow) was measured from the onset of the stimulus to the onset of the downward deflection of the earliest component of the field potential. The size of the evoked potentials (gray shading) was measured by the peak area under the baseline from the onset to the offset of the averaged waveform. The peak onset was defined as the time at which the waveform following the stimulation pulse crossed 2 SDs below the average of the baseline (dotted horizontal line), and the peak offset was vice versa (black arrow). Baseline (horizontal lines) were determined from the mean of the background activity (from 50 to 20 ms before stimulation). SR Stim, Time of stimulation to the SR. **B,** Example of DR-evoked LFP and MUA. The size of the DR-evoked potentials was measured by the peak area under the baseline from the onset to the negative peak of the averaged waveform. **C,** Example of SR-evoked incoming afferent volley recorded in the radial nerve cuff electrode (average of 2328 stim.). **D,** Example of DR-evoked incoming afferent volley recorded in the radial nerve cuff electrode (average of 2079 stim.). **E,** Boxplots showing the amplitude of the SR incoming volleys for each behavioral period (left; wrist flexion, right; wrist extension). Symbols denote data for the individual responses. Red crosses represent outlier data. One-way ANOVA with post hoc Sidak’s test; [“Flexion” (F_2,224_ = 0.38, p = 0.68), “Extension” (F_2,145_ = 0.126, p = 0.88)]. Note that no significant difference was found for incoming volleys among behavioral epochs in the SR stimulation condition. **F,** Boxplots showing amplitude of the DR incoming volleys for each behavioral period (left; wrist flexion, right; wrist extension). One-way ANOVA with post hoc Sidak’s test; [“Flexion” (F_2,92_ = 0.33, p = 0.71), “Extension” (F_2,93_ = 0.34, p = 0.71)]. Note that no significant difference was found for incoming volleys among behavioral epochs in the DR stimulation condition.

**Figure S3:**
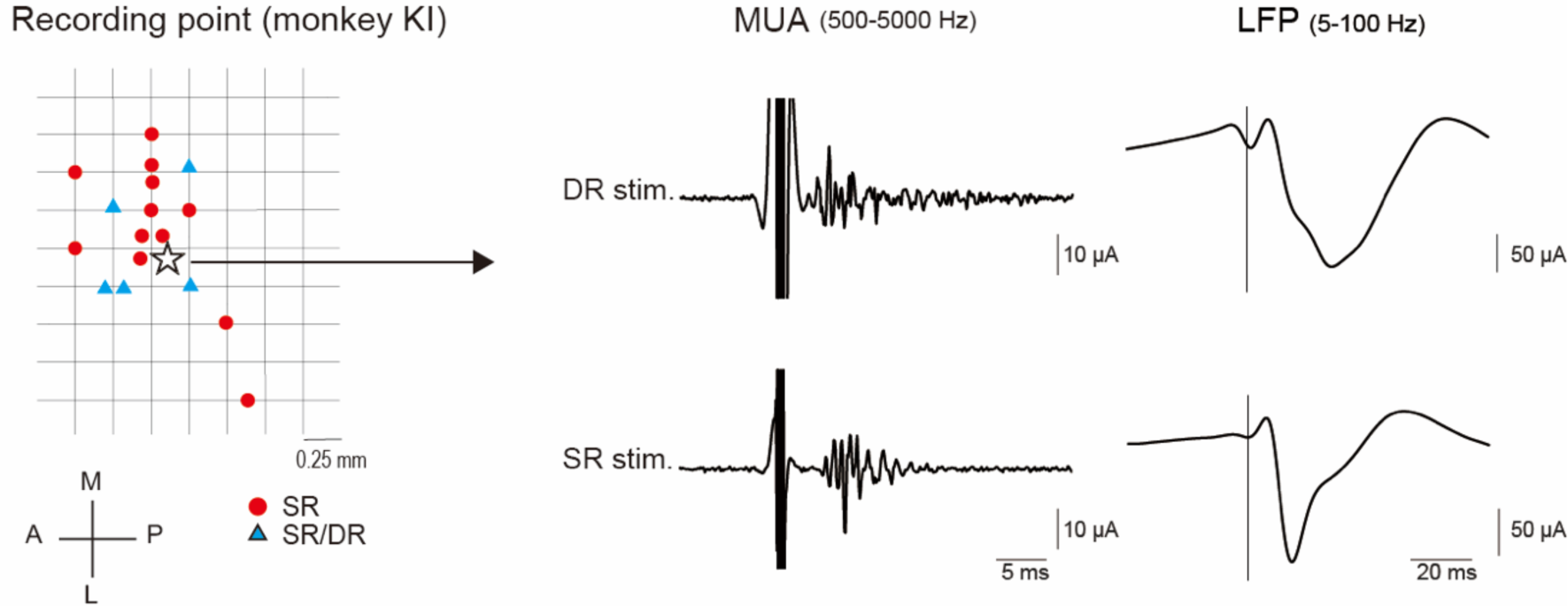
Typical example of multi-modal responses evoked by SR and DR nerve stimulation. **Left;** Electrode insert point map in monkey KI, same as ED Fig.1B. Star indicates the recording point, which shows in the right panel. **Right;** Averaged DR-(**Top**) and SR-(**Bottom**) evoked multi-unit activities (MUA, **Left**) and local field potentials (LFP, **Right**). The vertical line in right panels indicates the timing of nerve stimulation. Note that there is convergence in both cutaneous and proprioceptive afferent signals on this recording point.

**Figure S4:**
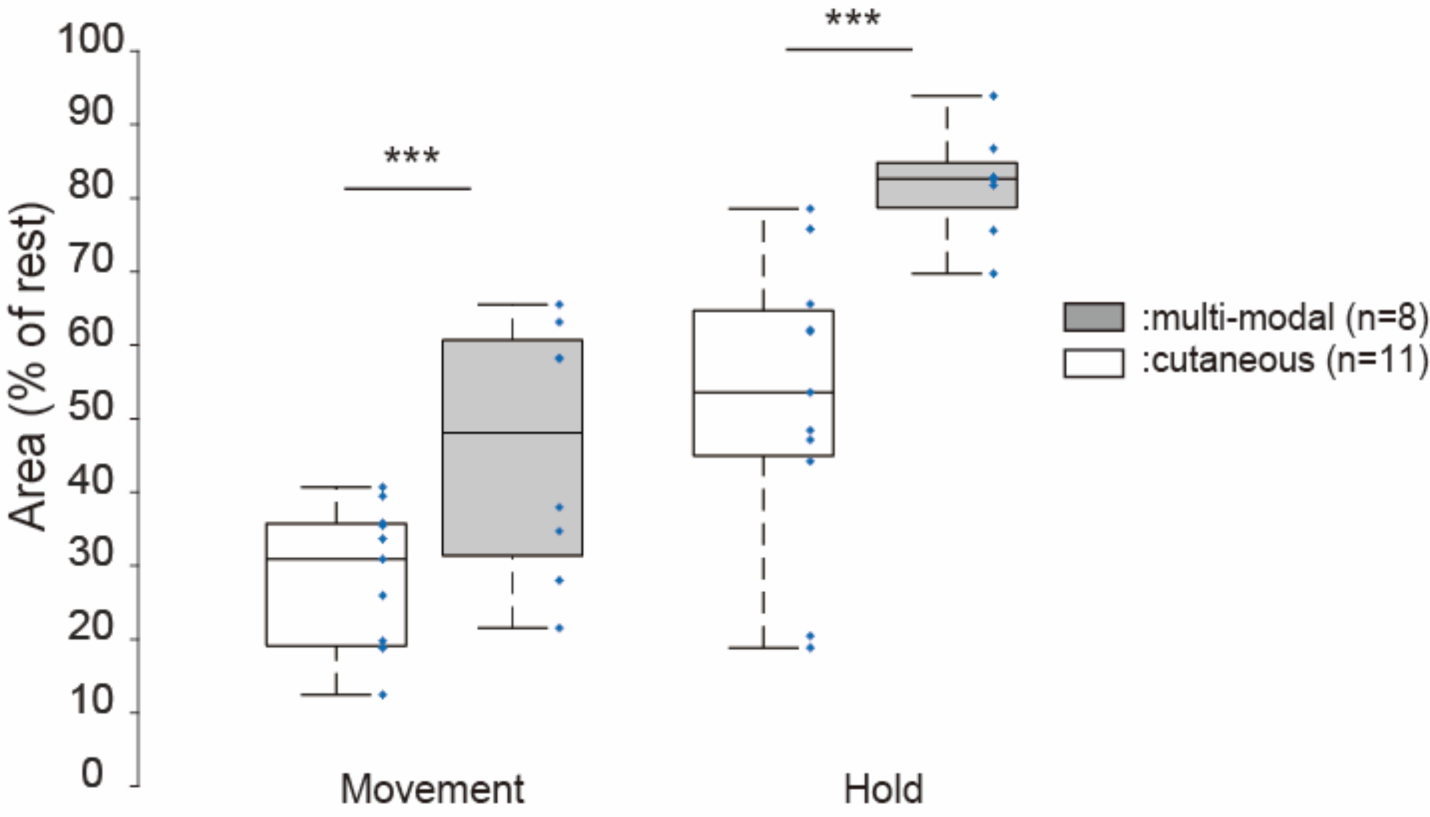
Comparison of SR-evoked local field potentials between the cutaneous and multi-modal area in the cuneate nucleus. Boxplot showing SR-evoked local field potentials (LFPs) recorded in the cutaneous (white) or the multi-modal (gray) area during voluntary wrist movement. The size of SR-evoked LFPs was combined regardless of movement direction and then pooled at each task epoch and normalized by the rest epoch. Symbols denote data for each recording sites. Two-way ANOVA with post hoc Sidak’s test; [“area” (F_1,34_ =24.148, p < 0.001), “epoch” (F_1,34_ = 39.313, p < 0.001)]: *** p < 0.001.

**Figure S5:**
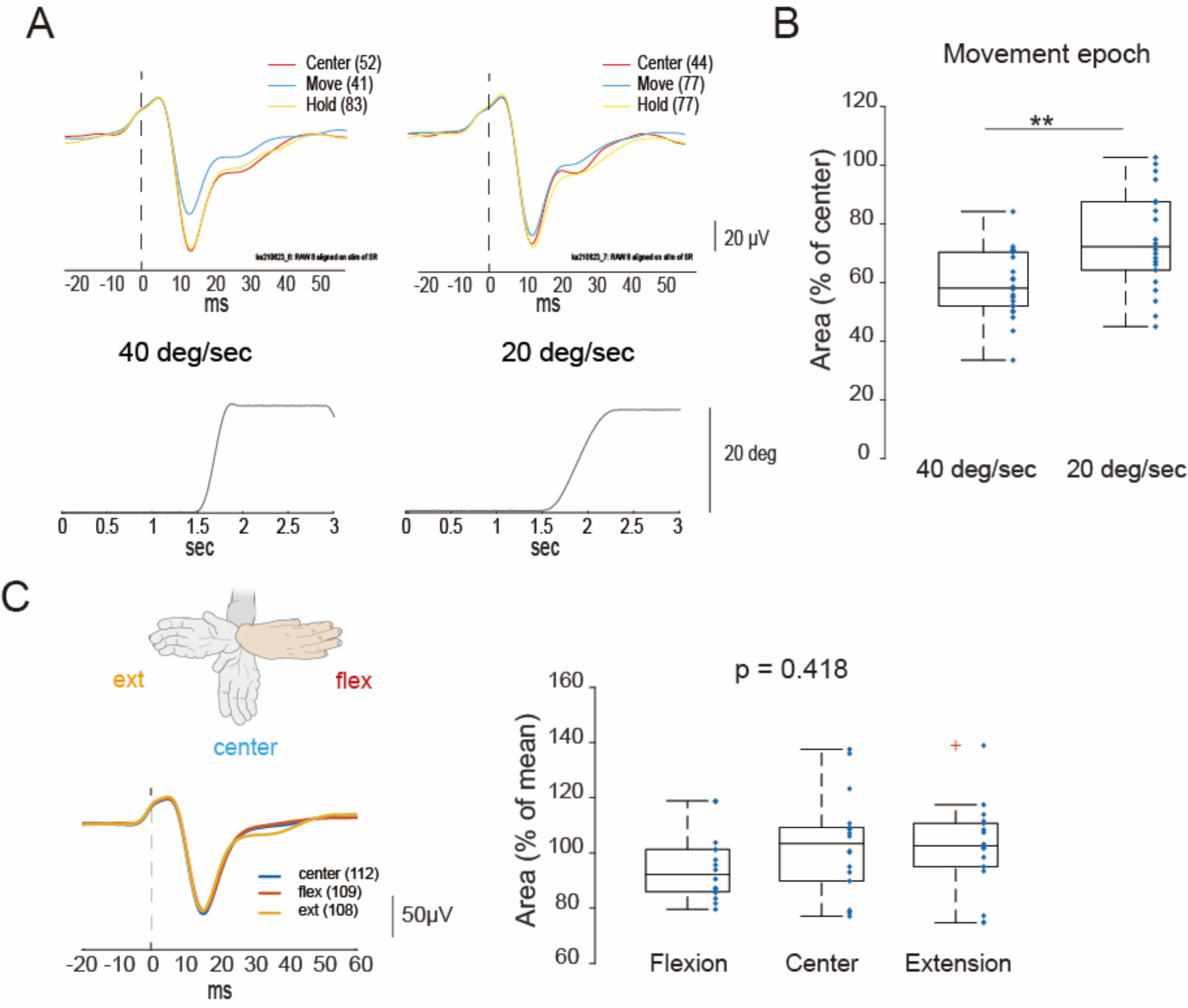
Modulation of SR-evoked local field potentials on the different movement speeds during passive movement. **A,** SR-evoked local field potentials (LFPs) (**Top**) and the displacement of wrist joint **(Bottom)** during passive wrist flexion. The manipulandum was automatically moved at 40 degrees/sec speed (**Left**) and 20 degrees/sec (**Right**), at a range of 20 degrees from the neutral position. Numbers in parentheses indicate the number of stimulation pulses used for averaging in a given behavioral epoch. **B,** Boxplots showing the size of SR-evoked LFPs during movement epoch at each movement speed condition. Symbols denote data for each recording site (n=22). Note that the size of evoked LFPs was more suppressed in fast speed conditions than slow speed. **significant difference between 40 deg/sec vs. 20 deg/sec, p=0.004: paired t-test. **C, Left;** Averaged SR-evoked LFPs in the center position (blue), flexion position (red; 20 degree flexed from the center), and extension position (orange; 20 degrees extended from the center). Numbers in parentheses indicate the number of stimulation pulses used for averaging in a given position. **Right;** Boxplots showing the size of SR-evoked LFPs that were obtained in each wrist position. Symbols denote data for each recording sites (n=18). Crosses represent outlier data. Note that the size of SR-evoked LFPs is not significantly different among wrist positions. One-way ANOVA with post hoc Sidak’s test; [“position” (F_2,51_ = 0.88, p = 0.418)].

**Figure S6:**
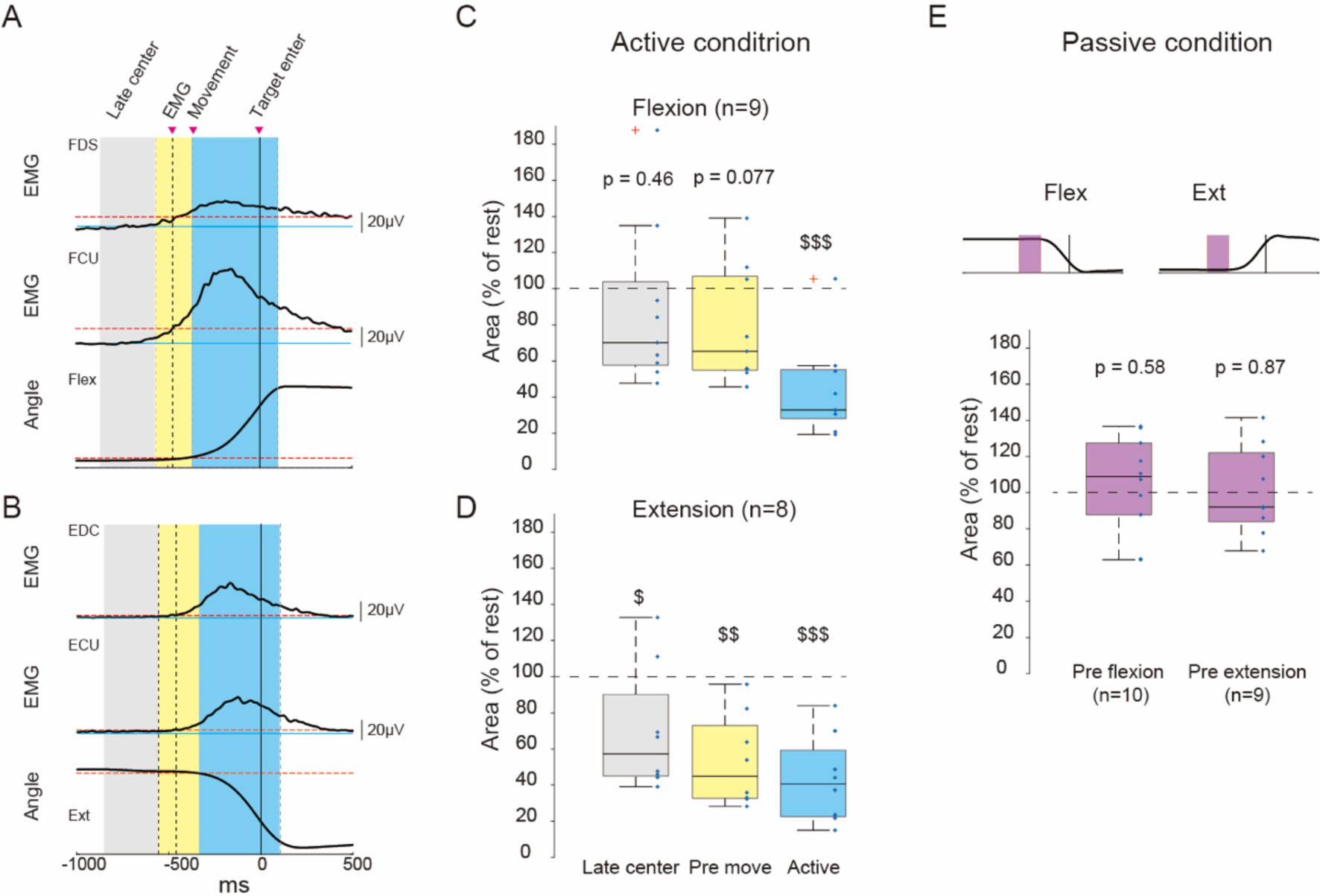
Modulation of SR-evoked local field potentials on before and during active movement. **A and B**, Example of the electromyogram (EMG) activities on FDS, FCU, EDC, ECU muscles and movement trajectory of wrist joint during active flexion (A) and extension (B) movement. EMGs and wrist angle were averaged within the session. The horizontal solid blue line indicates the mean value of EMGs and wrist angle during 500 ms preceding EMG onset. The horizontal dashed red line illustrates 4 standard deviations (SD) values above or below (in the angle of wrist extension) the mean. The timing at which the value of EMG and wrist angle crosses the dashed red line was defined as the EMG and movement onset point, respectively. The filled gray area represents a late center period, which ranged from 400 ms to 100 ms before EMG onset. The filled yellow area represents a pre-movement period, from 100 ms before EMG onset to movement onset. The blue shadow area represents the active movement period, from movement onset to movement offset point. Note that EMG onset was earlier than movement onset, whenever a wrist moved. **C and D,** Boxplots showing the size of SR evoked local field potentials (LFPs) that obtained late center, during the pre-movement and active flexion (C) and extension (D) movement period. Symbols denote data for each recording session. Red crosses represent outlier data. Note that the size of evoked potentials at the pre-movement period was suppressed in the extension movement direction. $ p < 0.05, $$ p < 0.01, $$$ p < 0.001 significant difference compared with the rest by one-sample t-test. **E,** Boxplots showing the size of SR evoked LFPs that were obtained during pre-movement in the passive condition. The pre-movement period was defied for 200 ms before the movement onset. Note that in the passive condition, there is no significant suppression on the size of evoked LFPs at the pre-movement period: pre-flexion, p = 0.583, pre-extension, p = 0.879: one-sample t-test.

**Figure S7:**
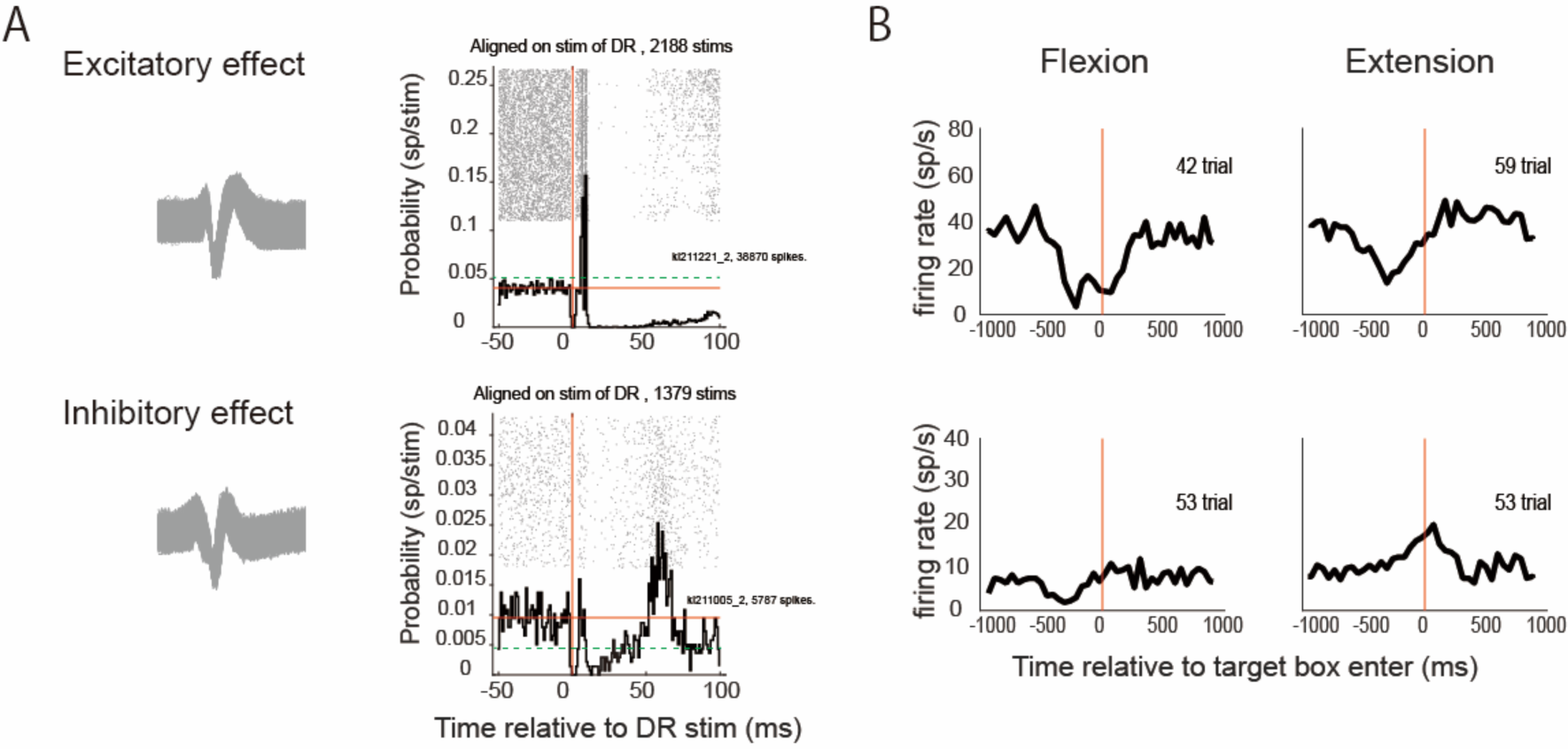
Response patterns of cuneate neurons elicited from DR nerve stimulation. **A,** Examples of the excitatory (**Top**) or inhibitory (**Bottom**) response of cuneate neuron, elicited from DR nerve stimulation. Cell discharge raster plot and its peristimulus time histogram (PSTH) of a cuneate neuron aligned on DR stimulation pulses. All formats are similar to Figure 4A. **B,** Averaged firing patterns of DR response neuron (shown in A) during a wrist flexion/extension task. Excitatory response neurons showed a decrease in firing rate in both flexion and extension movement, compared with the center epoch (p<0.001 in flexion and extension movement epoch, paired t-test). Inhibitory response neuron represents a decrease in its activity in flexion movement but an increase in extension movement, compared with the center epoch (p=0.009 in flexion, p<0.001 in extension, paired t-test). All formats are similar to Figure 4B.

## Reference

1. Johansson, R.S., and Flanagan, J.R. (2009). Coding and use of tactile signals from the fingertips in object manipulation tasks. Nature Reviews Neuroscience 10, 345–359. 10.1038/nrn2621.

2. Miall, R.C., Weir, D.J., and Stein, J.F. (1985). Visuomotor tracking with delayed visual feedback. Neuroscience 16, 511–520. 10.1016/0306-4522(85)90189-7.

3. Crevecoeur, F., McIntyre, J., Thonnard, J.L., and Lefèvre, P. (2010). Movement stability under uncertain internal models of dynamics. J Neurophysiol 104, 1301–1313. 10.1152/jn.00315.2010.

4. Papakostopoulos, D., Cooper, R., and Crow, H.J. (1975). Inhibition of cortical evoked potentials and sensation by self-initiated movement in man. Nature 258, 321–324. 10.1038/258321a0.

5. Rushton, D.N., Rothwell, J.C., and Craggs, M.D. (1981). Gating of somatosensory evoked potentials during different kinds of movement in man. Brain 104, 465–491. 10.1093/brain/104.3.465.

6. Cohen, L.G., and Starr, A. (1987). Localization, timing and specificity of gating of somatosensory evoked potentials during active movement in man. Brain 110 (Pt 2), 451–467. 10.1093/brain/110.2.451.

7. Blakemore, S.J., Wolpert, D.M., and Frith, C.D. (1998). Central cancellation of self-produced tickle sensation. Nat Neurosci 1, 635–640. 10.1038/2870.

8. Azim, E., and Seki, K. (2019). Gain control in the sensorimotor system. Curr Opin Physiol 8, 177–187. 10.1016/j.cophys.2019.03.005.

9. Kim, O.A., Forrence, A.D., and McDougle, S.D. (2022). Motor learning without movement. Proc Natl Acad Sci U S A 119, e2204379119. 10.1073/pnas.2204379119.

10. Gibson, J.J. (1962). Observations on active touch. Psychol Rev 69, 477–491. 10.1037/h0046962.

11. Wolpert, D.M., and Miall, R.C. (1996). Forward Models for Physiological Motor Control. Neural Netw 9, 1265–1279. 10.1016/s0893-6080(96)00035-4.

12. von Holst, E., and Mittelstaedt, H. (1950). The reafference principle: interaction between the central nervous system and the periphery. Die Naturwissenschaften 37, 464–476.

13. Sperry, R.W. (1950). Neural basis of the spontaneous optokinetic response produced by visual inversion. J Comp Physiol Psychol 43, 482–489. 10.1037/h0055479.

14. Miall, R.C., Weir, D.J., Wolpert, D.M., and Stein, J.F. (1993). Is the cerebellum a smith predictor? J Mot Behav 25, 203–216. 10.1080/00222895.1993.9942050.

15. Marshall, W.H. (1941). OBSERVATIONS ON SUBCORTICAL SOMATIC SENSORY MECHANISMS OF CATS UNDER NEMBUTAL ANESTHESIA. Journal of Neurophysiology 4, 25–43. 10.1152/jn.1941.4.1.25.

16. Coulter, J.D. (1974). Sensory transmission through lemniscal pathway during voluntary movement in the cat. J Neurophysiol 37, 831–845. 10.1152/jn.1974.37.5.831.

17. Chapman, C.E., Jiang, W., and Lamarre, Y. (1988). Modulation of lemniscal input during conditioned arm movements in the monkey. Experimental Brain Research 72, 316–334. 10.1007/BF00250254.

18. Seki, K., and Fetz, E.E. (2012). Gating of Sensory Input at Spinal and Cortical Levels during Preparation and Execution of Voluntary Movement. The Journal of Neuroscience 32, 890–902. 10.1523/jneurosci.4958-11.2012.

19. Seki, K., Perlmutter, S.I., and Fetz, E.E. (2003). Sensory input to primate spinal cord is presynaptically inhibited during voluntary movement. Nat Neurosci 6, 1309–1316. 10.1038/nn1154.

20. Confais, J., Kim, G., Tomatsu, S., Takei, T., and Seki, K. (2017). Nerve-Specific Input Modulation to Spinal Neurons during a Motor Task in the Monkey. The Journal of Neuroscience 37, 2612–2626. 10.1523/jneurosci.2561-16.2017.

21. Abraira, V.E., and Ginty, D.D. (2013). The sensory neurons of touch. Neuron 79, 618–639. 10.1016/j.neuron.2013.07.051.

22. Andersen, P., Eccles, J.C., Oshima, T., and Schmidt, R.F. (1964). MECHANISMS OF SYNAPTIC TRANSMISSION IN THE CUNEATE NUCLEUS. J Neurophysiol 27, 1096–1116. 10.1152/jn.1964.27.6.1096.

23. Andersen, P., Eccles, J.C., Schmidt, R.F., and Yokota, T. (1964). IDENTIFICATION OF RELAY CELLS AND INTERNEURONS IN THE CUNEATE NUCLEUS. J Neurophysiol 27, 1080–1095. 10.1152/jn.1964.27.6.1080.

24. Harris, F., Jabbur, S.J., Morse, R.W., and Towe, A.L. (1965). Influence of the Cerebral Cortex on the Cuneate Nucleus of the Monkey. Nature 208, 1215–1216. 10.1038/2081215a0.

25. Walberg, F. (1965). Axoaxonic contacts in the cuneate nucleus, probable basis for presynaptic depolarization. Exp Neurol 13, 218–231. 10.1016/0014-4886(65)90111-1.

26. Cole, J.D., and Gordon, G. (1983). Timing of corticofugal actions on the gracile and cuneate nuclei of the cat. J Physiol 341, 139–152. 10.1113/jphysiol.1983.sp014797.

27. Cheema, S., Whitsel, B.L., and Rustioni, A. (1983). The corticocuneate pathway in the cat: relations among terminal distribution patterns, cytoarchitecture, and single neuron functional properties. Somatosens Res 1, 169–205. 10.3109/07367228309144547.

28. Bentivoglio, M., and Rustioni, A. (1986). Corticospinal neurons with branching axons to the dorsal column nuclei in the monkey. J Comp Neurol 253, 260–276. 10.1002/cne.902530212.

29. Conner, J.M., Bohannon, A., Igarashi, M., Taniguchi, J., Baltar, N., and Azim, E. (2021). Modulation of tactile feedback for the execution of dexterous movement. Science 374, 316–323. doi:10.1126/science.abh1123.

30. Ghez, C., and Pisa, M. (1972). Inhibition of afferent transmission in cuneate nucleus during voluntary movement in the cat. Brain Res 40, 145–155. 10.1016/0006-8993(72)90120-5.

31. Richardson, A.G., Weigand, P.K., Sritharan, S.Y., and Lucas, T.H. (2016). A chronic neural interface to the macaque dorsal column nuclei. J Neurophysiol 115, 2255–2264. 10.1152/jn.01083.2015.

32. Suresh, A.K., Winberry, J.E., Versteeg, C., Chowdhury, R., Tomlinson, T., Rosenow, J.M., Miller, L.E., and Bensmaia, S.J. (2017). Methodological considerations for a chronic neural interface with the cuneate nucleus of macaques. J Neurophysiol 118, 3271–3281. 10.1152/jn.00436.2017.

33. Goodwin, A.W., Macefield, V.G., and Bisley, J.W. (1997). Encoding of object curvature by tactile afferents from human fingers. J Neurophysiol 78, 2881–2888. 10.1152/jn.1997.78.6.2881.

34. LaMotte, R.H., and Srinivasan, M.A. (1987). Tactile discrimination of shape: responses of rapidly adapting mechanoreceptive afferents to a step stroked across the monkey fingerpad. J Neurosci 7, 1672–1681. 10.1523/jneurosci.07-06-01672.1987.

35. LaMotte, R.H., and Srinivasan, M.A. (1987). Tactile discrimination of shape: responses of slowly adapting mechanoreceptor afferents to a step stroked across the monkey fingerpad. J Neurosci 7, 1655–1671. 10.1523/jneurosci.07-06-01655.1987.

36. Berkley, K.J., Budell, R.J., Blomqvist, A., and Bull, M. (1986). Output systems of the dorsal column nuclei in the cat. Brain Research Reviews 11, 199–225. https://doi.org/10.1016/0165-0173(86)90012-3.

37. Loutit, A.J., Vickery, R.M., and Potas, J.R. (2021). Functional organization and connectivity of the dorsal column nuclei complex reveals a sensorimotor integration and distribution hub. J Comp Neurol 529, 187–220. 10.1002/cne.24942.

38. Miocinovic, S., Zhang, J., Xu, W., Russo, G.S., Vitek, J.L., and McIntyre, C.C. (2007). Stereotactic neurosurgical planning, recording, and visualization for deep brain stimulation in non-human primates. Journal of neuroscience methods 162, 32–41. 10.1016/j.jneumeth.2006.12.007.

39. Seki, K., Perlmutter, S.I., and Fetz, E.E. (2009). Task-dependent modulation of primary afferent depolarization in cervical spinal cord of monkeys performing an instructed delay task. J Neurophysiol 102, 85–99. 10.1152/jn.91113.2008.

40. Florence, S.L., Wall, J.T., and Kaas, J.H. (1989). Somatotopic organization of inputs from the hand to the spinal gray and cuneate nucleus of monkeys with observations on the cuneate nucleus of humans. J Comp Neurol 286, 48–70. 10.1002/cne.902860104.

41. Jiang, W., Lamarre, Y., and Chapman, C.E. (1990). Modulation of cutaneous cortical evoked potentials during isometric and isotonic contractions in the monkey. Brain Res 536, 69–78. 10.1016/0006-8993(90)90010-9.

42. Greenspan, J.D. (1992). Influence of velocity and direction of surface-parallel cutaneous stimuli on responses of mechanoreceptors in feline hairy skin. J Neurophysiol 68, 876–889. 10.1152/jn.1992.68.3.876.

43. Edin, B., Essick, G., Trulsson, M., and Olsson, K. (1995). Receptor encoding of moving tactile stimuli in humans. I. Temporal pattern of discharge of individual low-threshold mechanoreceptors. The Journal of Neuroscience 15, 830–847. 10.1523/jneurosci.15-01-00830.1995.

44. Cheney, P.D., and Preston, J.B. (1976). Classification and response characteristics of muscle spindle afferents in the primate. J Neurophysiol 39, 1–8. 10.1152/jn.1976.39.1.1.

45. Ferrington, D.G., Rowe, M.J., and Tarvin, R.P. (1986). High gain transmission of single impulses through dorsal column nuclei of the cat. Neurosci Lett 65, 277–282. 10.1016/0304-3940(86)90274-0.

46. Iwamura, Y. (1998). Hierarchical somatosensory processing. Curr Opin Neurobiol 8, 522–528. 10.1016/s0959-4388(98)80041-x.

47. Sobinov, A.R., and Bensmaia, S.J. (2021). The neural mechanisms of manual dexterity. Nat Rev Neurosci 22, 741–757. 10.1038/s41583-021-00528-7.

48. Florence, S.L., Wall, J.T., and Kaas, J.H. (1991). Central projections from the skin of the hand in squirrel monkeys. Journal of Comparative Neurology 311, 563–578. https://doi.org/10.1002/cne.903110410.

49. Versteeg, C., Chowdhury, R.H., and Miller, L.E. (2021). Cuneate nucleus: the somatosensory gateway to the brain. Current Opinion in Physiology 20, 206–215. https://doi.org/10.1016/j.cophys.2021.02.004.

50. Pruszynski, J.A., Flanagan, J.R., and Johansson, R.S. (2018). Fast and accurate edge orientation processing during object manipulation. Elife 7. 10.7554/eLife.31200.

51. Jones, E.G. (2000). Cortical and subcortical contributions to activity-dependent plasticity in primate somatosensory cortex. Annu Rev Neurosci 23, 1–37. 10.1146/annurev.neuro.23.1.1.

52. Edin, B.B., and Abbs, J.H. (1991). Finger movement responses of cutaneous mechanoreceptors in the dorsal skin of the human hand. J Neurophysiol 65, 657–670. 10.1152/jn.1991.65.3.657.

53. Edin, B.B., and Vallbo, A.B. (1990). Dynamic response of human muscle spindle afferents to stretch. J Neurophysiol 63, 1297–1306. 10.1152/jn.1990.63.6.1297.

54. Kakuda, N., Vallbo, A.B., and Wessberg, J. (1996). Fusimotor and skeletomotor activities are increased with precision finger movement in man. J Physiol 492 (Pt 3), 921–929. 10.1113/jphysiol.1996.sp021358.

55. Chirila, A.M., Rankin, G., Tseng, S.Y., Emanuel, A.J., Chavez-Martinez, C.L., Zhang, D., Harvey, C.D., and Ginty, D.D. (2022). Mechanoreceptor signal convergence and transformation in the dorsal horn flexibly shape a diversity of outputs to the brain. Cell 185, 4541–4559.e4523. 10.1016/j.cell.2022.10.012.

56. Witham, C.L., and Baker, S.N. (2011). Modulation and transmission of peripheral inputs in monkey cuneate and external cuneate nuclei. J Neurophysiol 106, 2764–2775. 10.1152/jn.00449.2011.

57. Jörntell, H., Bengtsson, F., Geborek, P., Spanne, A., Terekhov, A.V., and Hayward, V. (2014). Segregation of tactile input features in neurons of the cuneate nucleus. Neuron 83, 1444–1452. 10.1016/j.neuron.2014.07.038.

58. Suresh, A.K., Greenspon, C.M., He, Q., Rosenow, J.M., Miller, L.E., and Bensmaia, S.J. (2021). Sensory computations in the cuneate nucleus of macaques. Proceedings of the National Academy of Sciences 118, e2115772118. doi:10.1073/pnas.2115772118.

59. Doyle, M.W., and Andresen, M.C. (2001). Reliability of monosynaptic sensory transmission in brain stem neurons in vitro. J Neurophysiol 85, 2213–2223. 10.1152/jn.2001.85.5.2213.

60. Andersen, P., Eccles, J.C., Schmidt, R.F., and Yokota, T. (1964). DEPOLARIZATION OF PRESYNAPTIC FIBERS IN THE CUNEATE NUCLEUS. J Neurophysiol 27, 92–106. 10.1152/jn.1964.27.1.92.

61. Jabbur, S.J., and Banna, N.R. (1968). Presynaptic inhibition of cuneate transmission by widespread cutaneous inputs. Brain Res 10, 273–276. 10.1016/0006-8993(68)90135-2.

62. Biedenbach, M.A., Jabbur, S.J., and Towe, A.L. (1971). Afferent inhibition in the cuneate nucleus of the rhesus monkey. Brain Research 27, 179–183. https://doi.org/10.1016/0006-8993(71)90381-7.

63. Drucker, C.B., Carlson, M.L., Toda, K., DeWind, N.K., and Platt, M.L. (2015). Non-invasive primate head restraint using thermoplastic masks. Journal of neuroscience methods 253, 90–100. 10.1016/j.jneumeth.2015.06.013.

